# Dual curvature sensing governs cell orientation and curvotaxis

**DOI:** 10.64898/2026.05.09.723774

**Authors:** Shao-Zhen Lin, Kosei Tomida, Boon Heng Ng, Chin Hao Lee, Juin Shin Wee, Zixuan Zhao, Eliza Li Shan Fong, Chii Jou Chan

**Author notes:** These authors contributed equally to this work.

## Abstract

Cells lying in a curved environment can respond to the surface curvature by reorienting their shape. However, whether cells respond to the mean curvature and/or the Gaussian curvature remains largely unexplored. Here, inspired by experimental observations of how ovarian theca cells (TCs) orient themselves on substrates with different curvatures, we propose a theoretical framework for active nematic layers on curved surfaces. In this model, we assume that the nematic directors of the cells respond to both the mean curvature and the Gaussian curvature of the underlying substrate surface. Our theory predicts specific cell orientation patterns on hemicylindrical, hourglass- and dome-like substrates, consistent with experimental observations. In addition, by incorporating a curvotaxis traction, our model successfully recapitulates the experimental observation of TC accumulation at convex regions of hemicylindrical substrates as well as saddle-shaped regions of more complex geometries. Overall, our work reveals the unexpected role of cell curvature sensing in driving collective migration and pattern formation on various substrate curvatures.

**SIGNIFICANCE:** Substrate surface curvature is a critical environmental cue that can influence multicellular organization and functions. Yet how cells collectively align and migrate on complex curved surfaces remains unclear. Here, we proposed a hydrodynamic theory of active nematic layers over curved surfaces for contractile theca cells (TCs), where we assume that the nematic directors of cells can respond to both the mean curvature and the Gaussian curvature of the underlying substrates. Our theory predicts distinct cell orientation patterns on hemicylindrical, hourglass- and dome-like substrates, consistent with experimental observations. Furthermore, by introducing curvotaxis traction, our model recapitulates experimentally observed accumulation of TCs at the convex regions of hemicylindrical substrates as well as saddle-shaped regions of more complex geometries. Together, our study provides a simple theoretical framework to unify our understanding of curvature sensing across complex topology, providing insights into geometric control of tissue pattern formation.

## INTRODUCTION

Cell orientation and movement are fundamental biological processes that play critical roles in tissue development, wound healing, and disease progression. These behaviors are tightly regulated by both biochemical and biophysical cues from the extracellular environment, with emerging evidence highlighting the profound influence of surface curvature (of the substrate where cells adhere to) [1–3].

Cells can sense the surface curvature and reorient their shape along the directions of either maximal or minimal curvature [1]. For example, when adhering to the outer surface of a cylinder, epithelial cells (e.g., Madin-Darby canine kidney (MDCK) cells [4–8]) tend to orient their shape along the circumferential direction (i.e., the maximal curvature direction), whereas fibroblasts tend to orient along the long axis direction (i.e., the minimal curvature direction) [1, 9]. While previous studies have shed light on cellular response to simple surfaces, e.g., sinusoidal surfaces [10, 11], cylinders [4, 5, 8, 12], and spheres [13–17], it remains unclear how cells can sense and distinguish the distinct shape (mean curvature) and topology (Gaussian curvature) of complex surface geometries [2]. Interestingly, a recent work has highlighted the importance of Gaussian curvature in patterning nematic order and driving emergent tissue shapes in programmable living materials [18].

On the theory side, to describe cell orientation in response to the surface curvature, previous studies proposed a coupling between cell orientation and surface curvature in an effective free energy formalism [19–22]. To describe the up-down (apical-basal) asymmetry of cells adhering to a substrate surface, in the linear order, such a curvature-sensing free energy can be expressed as [22] (Supporting Information, Sec. I):

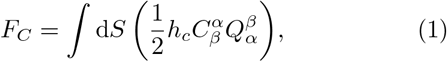

where 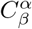 represents the extrinsic curvature tensor of the substrate surface [22, 23] and 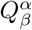 is a nematic tensor that describes cell shape and orientation (Fig. 2(b)). In Eq. (1), *h*_*c*_ is a constant parameter describing how a cell orients its shape in response to surface curvature, thus referred to as a curvature-sensing parameter: when *h*_*c*_ > 0, cells prefer to orient along the minimal curvature direction, e.g., the long axis direction of the outer surface of a cylinder; when *h*_*c*_ < 0, cells tend to orient along the maximal curvature direction, e.g., the circumferential direction of the outer surface of a cylinder. The curvature-sensing free energy Eq. (1) can describe the orientation feature of some cells over simple curved substrates, e.g., MDCK cells and fibroblasts over a cylindrical surface [1]. However, it remains unexplored whether it can be applied to other cell types and more complex curved geometries.

In this study, we explore the orientation response of primary ovarian theca cells (TCs) over various curved surfaces, and find that a constant parameter *h*_*c*_ in Eq. (1) cannot account for cell orientation on these surfaces. Instead, by incorporating a *h*_*c*_ that depends on both the mean curvature and the Gaussian curvature of the substrate surface into an active nematic gel theory, we successfully recapitulate the cell orientation behavior observed experimentally in various curved geometries. Furthermore, we demonstrate that a curvotaxis traction force, which depends on both the gradient of the mean curvature and the gradient of the Gaussian curvature, can account for the accumulation of TCs at certain curved surface regions. Together, our study reveals that cells can sense both the shape and topology of substrate curvature in their microenvironment and manifest collective nematic order and migration behavior accordingly.

## RESULTS

### *h*_*c*_ depends on substrate surface curvature

We cultured primary TCs over a hemicylindrical substrate (Fig. 1(a-c)); see Methods for details. We observed that, in the hill region (i.e., the convex region with a positive curvature), TCs exhibit collective orientation along the long axis of the hemicylindrical substrate (Fig. 1(a-c)), i.e., the minimal curvature direction. This indicates that *h*_*c*_ > 0. In contrast, in the valley region (i.e., the concave region with a negative curvature), TCs orient randomly without a preferred orientation direction (Fig. 1(a-c)), thus indicating that *h*_*c*_ ≈ 0. Furthermore, by placing hydrogel beads in touch with ovarian follicles, we were able to generate TC-coated bead doublets (see Methods for details). We observed that at the neck region (i.e., saddle-shaped region with a negative Gaussian curvature) of the bead doublet, TCs tend to orient along the circumferential direction (Fig. 1(d,e)), i.e., the maximal curvature direction. This indicates that *h*_*c*_ < 0, which is in sharp contrast to the orientation behavior of TCs at the hill region of a hemicylindrical substrate. This apparent contradiction (Fig. 1(f)) calls for a need to reconsider the curvature-sensing free energy Eq. (1) and suggests that a constant *h*_*c*_ can not account for all orientation responses of TCs on different substrate curvatures. One simple solution is to assume that *h*_*c*_ depends on the surface curvature 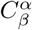, i.e., 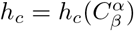.

**FIG. 1.**
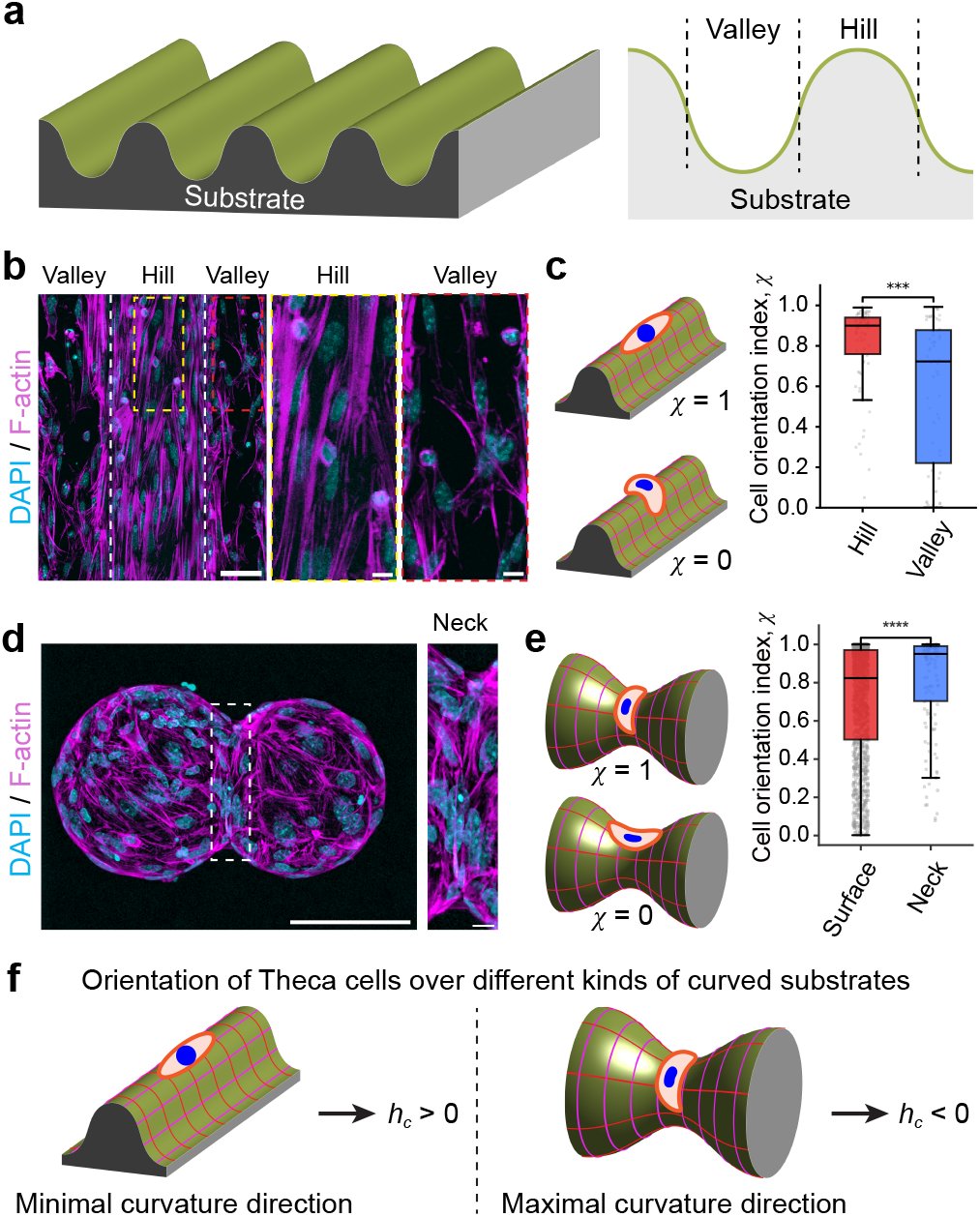
Orientation feature of TCs over curved substrates, including (a-c) a hemicylindrical substrate and (d,e) a bead doublet substrate. (a) Sketch of a hemicylindrical substrate. (b) Maximum projection of fluorescence images of TCs over a hemicylindrical substrate. Scale bar, 100 *µ*m. White boxes represent the zoomed-in hill region (left) and valley region (right). Scale bar, 10 *µ*m. (c) Left: Schematic showing cell orientation index of *χ* = 0 and 1. Right: Quantification of TC orientations at hill and valley regions on hemicylindrical substrates. Statistical significance was assessed using the two-sided Mann–Whitney U test. ^∗∗∗^*P* < 0.001; *n* = 74 nuclei for hill, *n* = 59 nuclei for valley; 3 regions from 3 independent experiments were analysed. (d) Maximum projection of fluorescence images of TCs over a bead doublet. Scale bar, 100 *µ*m. The white box represents the zoomed-in neck region. Scale bar, 10 *µ*m. (e) Left: Schematics of cell orientation index at *χ* = 0 and 1. Right: Quantification of TC orientation at the neck region. Statistical significance was assessed using the two-sided Mann–Whitney U test. ^∗∗∗∗^*P* < 0.0001; *n* = 1542 nuclei for surface, *n* = 134 nuclei for neck region; 10 doublets from 6 independent experiments were analysed. (f) Schematic showing different TC orientations on different curved substrates, suggesting an apparent contradiction.

**FIG. 2.**
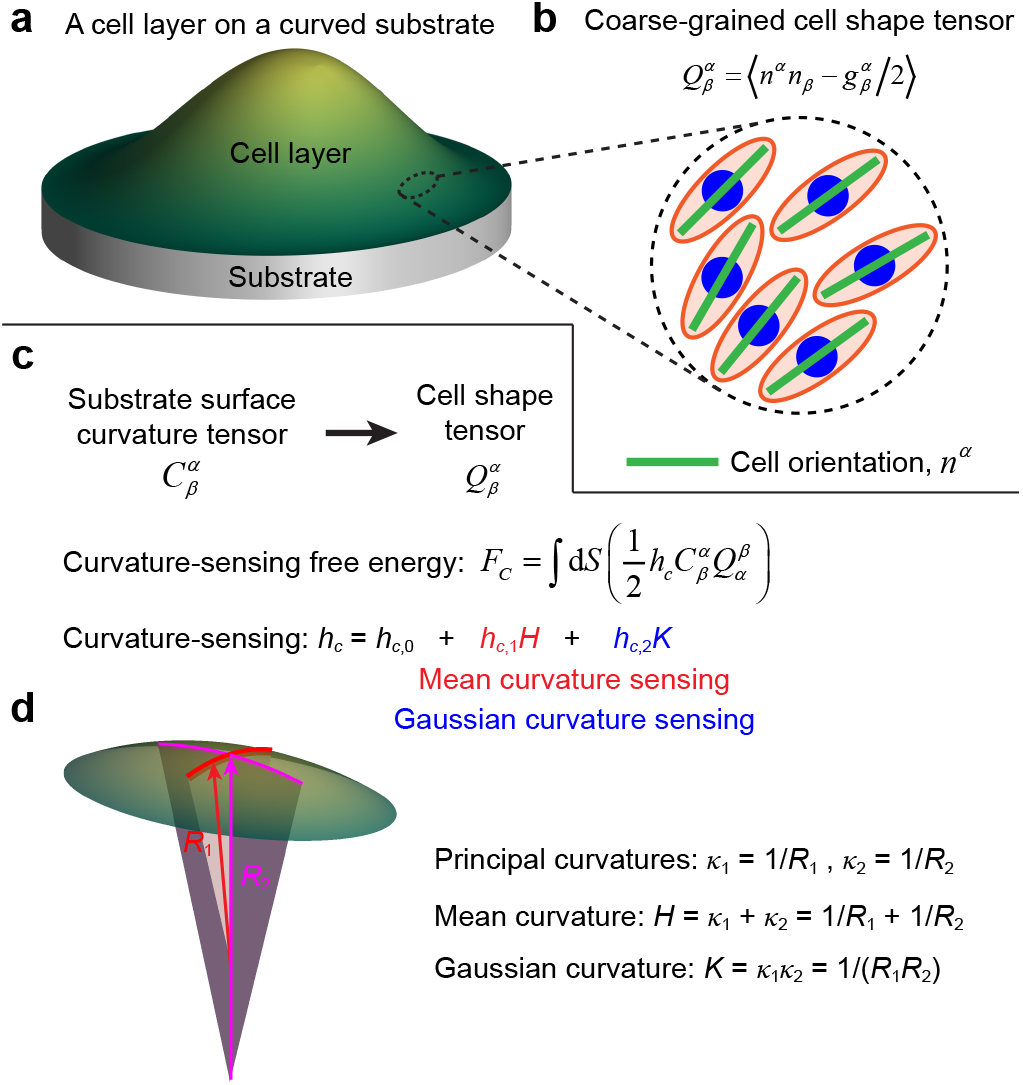
Schematic depicting the geometrical terms of curvature and the curvature-sensing free energy in the model. (a) A cell layer adhering on a curved substrate. (b) The spatial orientation of cells is described by a nematic tensor 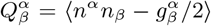, which corresponds to the locally average orientation of cells with *n*^*α*^ representing the orientation director of individual cells and ⟨·⟩ a local average. (c) The substrate curvature 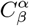 affects the orientation of cells by minimizing a curvature-sensing free energy *F*_*C*_ (see Eq. (1)) where the curvature-sensing parameter *h*_*c*_ could depend on the mean curvature 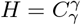 and the Gaussian curvature 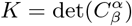. (d) Schematic of the principal curvatures *κ*_1_ and *κ*_2_, the mean curvature *H*, and the Gaussian curvature *K*.

### Active nematic gel theory for curved geometries

To explain our experimental observations, we propose an active nematic gel theory for curved geometries [22]. Let us consider a generic curved surface embedded in ℝ^3^ (Fig. 2(a)). Let ***r*** = ***r***(*x*^1^, *x*^2^) be the position on such a generic surface parametrized by the coordinates (*x*^1^, *x*^2^). Thus, the extrinsic curvature tensor is defined as *C*_*αβ*_ = −∂_*α*_∂_*β*_***r* · *m*** [22–25], where ***m*** = ***g***_1_ ×***g***_2_*/* |***g***_1_ ×***g***_2_| is the unit normal vector of the surface with ***g***_*α*_ = ∂***r****/*∂*x*^*α*^ being a basis of covariant vectors on the tangent plane. The dynamics of a compressible, active nematic layer adhering to a curved substrate can be characterized by three fields: (1) The nematic tensor 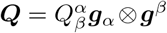, which describes the coarse-grained cell shape and orientation (Fig. 2(b)), thus also referred to as the cell shape tensor; (2) The flow vector ***v*** = *v*^*α*^***g***_*α*_, which describes the coarse-grained cell motion velocity; (3) The cell density *ρ*, which describes the average number of cells per unit area. We next give the detailed governing equations.

Following previous studies of active nematics [21, 26–29], we write the dynamic equation of 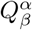 as:

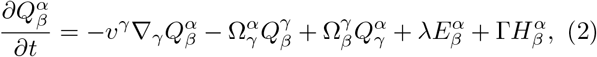

which considers the effects of advection, co-rotation, shear flow alignment, and relaxation to the energy-minimum state. In details, ∇_*γ*_ is the covariant derivative [21, 30, 31]; 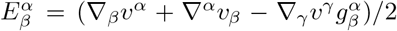 is the strain rate tensor; 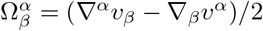 is the vorticity tensor; *λ* is a flow alignment coefficient; G describes the rate of cell shape relaxation to an energy-minimum state; 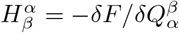 is a molecular field with *F* being the effective free energy of the system.

We write the effective free energy in two terms: *F* = *F*_0_ + *F*_*C*_. Here, *F*_0_ accounts for the Frank free energy of a nematic layer on a flat substrate and can be expressed as [22]: *F*_0_ = *f*_0_d*S*, where 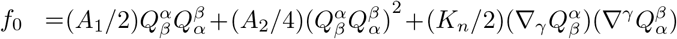 and 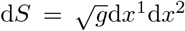 with *g* = det(*g*_*αβ*_) and *A*_1_ < 0, *A*_2_ > 0, and *K*_*n*_ > 0 being material constants. To describe the re-orientation of cells in response to the substrate surface curvature, we include the curvature-sensing free energy Eq. (1), see Fig. 2(c). Compared to previous studies [22], here, we assume that the curvature-sensing parameter *h*_*c*_ is not a constant, but instead, depends on the surface curvature 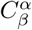, that is, 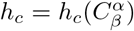. In the general case, we can write the Taylor expansion: 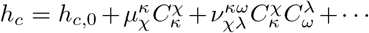, where 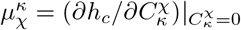 and 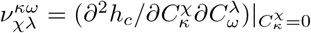. Considering both the mean curvature (denoted 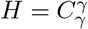) and the Gaussian curvature (denoted 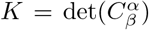) of the substrate surface, here, we further express the relation between *h*_*c*_ and 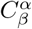 as the following (Fig. 2(c, d)):

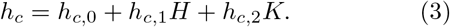

In Eq. (3), the first coefficient *h*_*c*,0_ quantifies how a cell aligns to a homogeneous curvature: *h*_*c*,0_ > 0 for cells that prefer to align along the minimal curvature direction, e.g., fibroblasts [1]; *h*_*c*,0_ < 0 for the opposite case, e.g., MDCK cells [1]. The second coefficient *h*_*c*,1_ accounts for the mean curvature-induced non-linear effect of cell re-orientation response [8]: *h*_*c*,1_ > 0 means that a positive mean curvature will enhance the re-orientation of cells along the minimal curvature direction. The third term *h*_*c*,2_ accounts for the Gaussian curvature-induced non-linear effect of cell re-orientation response. It should be noted that we have ignored the *H*^2^ term in Eq. (3), because such a quadratic term can not reflect the orientation symmetry breaking induced by the sign of the mean curvature *H*.

The flow velocity ***v*** satisfies the force balance equation:

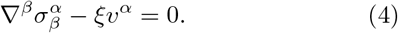

Here, *ξ* is the friction coefficient between cells and the substrate. The stress tensor 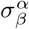 can be expressed as 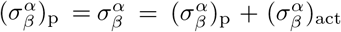 [26, 27, 32, 33], where 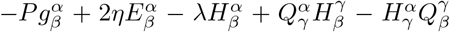 is the passive stress with *P* being the hydrodynamic pressure and *η* the viscosity; 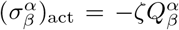 is the active stress. *ζ* quantifies cell activity: *ζ* < 0 for cells that actively contract along their shape direction, e.g., TCs in our experiments [34]; otherwise, *ζ* > 0, e.g., MDCK cells and retinal pigment epithelial cells [35]. In Eq. (4), we have ignored the inertial force because tissue flows usually take place in a low Reynolds number with a typical value ∼10^−5^ [36, 37]. We have also ignored active traction forces induced by splay and bend deformations of the nematic director field [22, 38] due to the lack of experimental evidence of its importance with TCs.

The cell density *ρ* satisfies the mass conservation equation:

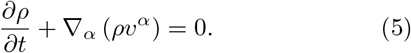

Here, we do not consider cell extrusions or divisions, which will be briefly discussed in the Discussion Section.

Finally, to close the above equations, one needs the constitutive equation for the pressure *P*, that is, to relate the pressure *P* to the three fields (***Q, v***, *ρ*). Following Ref. [39], here, we consider *P* = *P*_0_ ln(*ρ/ρ*_0_), where *P*_0_ is a reference pressure and *ρ*_0_ is a reference cell density.

### Deducing *h*_*c*,0_ and *h*_*c*,1_ from TC orientation on hemicylindrical substrates

We first apply our model to the hemicylindrical substrate, mimicked by a curved strip surface, which is curved only along one direction (denoted *s*) but possesses zero curvature along the perpendicular direction (denoted *y*); see Supporting Information, Sec. IIA.

While the effect of *h*_*c*,2_ is screened over the curved strip surface, the sign of *h*_*c*,0_ and *h*_*c*,1_ can be deduced by examining the cell orientation feature at regions of different curvatures. Specifically, in our experiments, we observed that TCs at the hill region (with a positive mean curvature, denoted *C*_hill_ > 0) orient well along the *y* axis (Fig. 1(a-c)), suggesting that *h*_*c*_(*C*_hill_) = *h*_*c*,0_ + *h*_*c*,1_*C*_hill_ > 0. In comparison, TCs at the valley region (with a negative mean curvature, denoted *C*_valley_ < 0) exhibit random orientation (Fig. 1(a-c)), suggesting *h*_*c*_(*C*_valley_) = *h*_*c*,0_ + *h*_*c*,1_*C*_valley_ ≈ 0. Therefore, combining these two equations results in *h*_*c*,0_ > 0 and *h*_*c*,1_ > 0 for TCs. We next perform numerical simulations to validate this inference. Indeed, with *h*_*c*,0_ > 0 and *h*_*c*,1_ > 0, our simulation agrees well with experiment, as shown in Fig. 3(a, b).

**FIG. 3.**
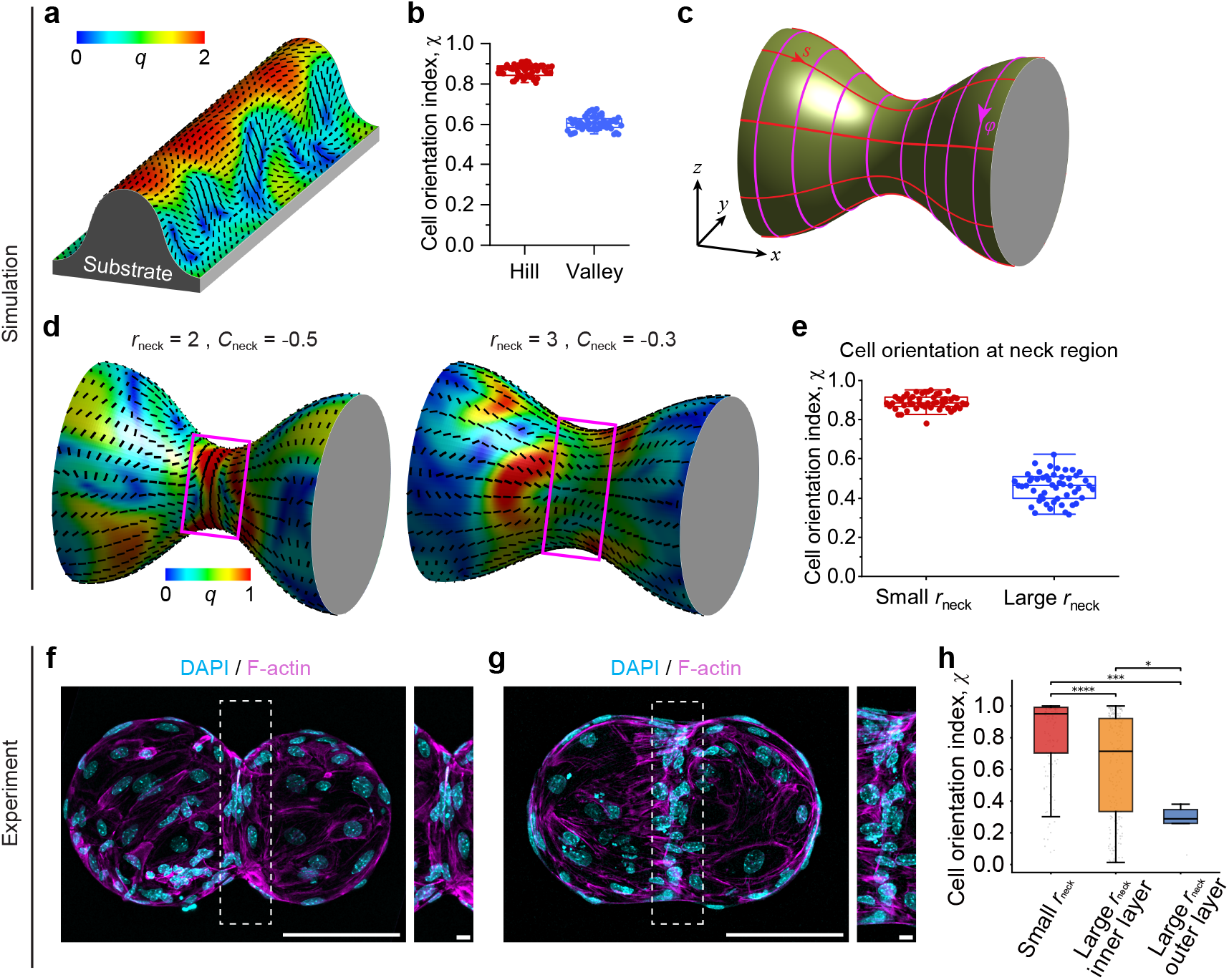
Numerical simulations of cell orientations based on a curvature-dependent *h*_*c*_. (a-e) Numerical simulations: (a, b) a curved strip geometry; (c-e) an hourglass geometry. (a) The cell elongation and orientation map over a curved strip substrate. The color code represents the magnitude of cell elongation *q*; the black lines indicate the local direction of cell orientation. (b) Comparison of the cell orientation index *χ* for cells at the hill region and for cells at the valley region. (c) Schematic of an hourglass geometry mimicking the neck region of the bead doublet in experiment: 3D view, along with the (*s, φ*) coordinates. (d) Comparison of the cell orientation maps at two different neck curvatures: (*top*) *r*_neck_ = 2 *µ*m and *C*_neck_ = −0.5, thus the mean curvature *H*_neck_ = 0 and the Gaussian curvature *K*_neck_ = −1; (*bottom*) *r*_neck_ = 3 *µ*m and *C*_neck_ = −0.3, thus the mean curvature *H*_neck_ = 0.03 and the Gaussian curvature *K*_neck_ = −0.9. (e) Comparison of cell orientation index *χ* at the neck region (see boxes in (d)) for a small *r*_neck_ and for a large *r*_neck_. See Supporting Information for parameter values. (f, g) Maximum projection of fluorescence images of TCs over a bead doublet with small (f) and large neck radius (g). Scale bar, 100 *µ*m. The white box represents zoomed-in neck regions. Scale bar, 10 *µ*m. (h) Quantification of TC orientation index at the neck region on bead doublets with small or large neck radius. Statistical significance was assessed using a two-sided Mann–Whitney U test. ^∗^*P* < 0.05,^∗∗∗^ *P* < 0.001,^∗∗∗∗^ *P* < 0.0001; *n* = 134 nuclei, *N* = 10 doublets from 6 experiments for small neck; *n* = 201 nuclei, *N* = 2 doublets from two experiments for large neck, inner layer; *n* = 5 nuclei, *N* = 2 doublets from two experiments for large neck, outer layer.

### Model recapitulates cell orientation behaviors over saddle-shaped regions of curved substrates

We next explore cell orientation features over complex surfaces of non-zero Gaussian curvature (including an hourglass and dome-like surface), focusing on saddle-shaped regions.

Let us first consider an hourglass surface, as shown in Fig. 3(c). We parameterize such a curved surface by the coordinates (*s, φ*) with *s* being the arc length coordinate (along the long axis of the hourglass) and *φ* the circumferential angle, see Fig. 3(c). For such a surface, using the coordinates (*s, φ*), we can simplify the governing equations (2), (4), and (5), expressed in terms of the nematic tensor components 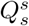 and 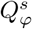, the flow velocity components *v*^*s*^ and *v*^*φ*^, and the cell density *ρ* (see Supporting Information, Sec. IIB).

We employ an hourglass geometry to mimic the neck region of the bead doublet in our experiment (Fig. 1(d,e)), by designing the curvature profile (see Supporting Information, Sec. IIB): at the neck region, the mean curvature *H*_neck_ = 1*/r*_neck_ + *C*_neck_ and the Gaussian curvature *K*_neck_ = *C*_neck_*/r*_neck_ with *r*_neck_ being the radius of the neck and *C*_neck_ the curvature along the long axis direction (i.e. *s* direction) at the neck. Note that *r*_neck_ > 0 and *C*_neck_ < 0, thus *K*_neck_ < 0; while the mean curvature *H*_neck_ can be either positive or negative, depending on the magnitudes of *r*_neck_ and *C*_neck_. The sign of *h*_*c*_ depends on *r*_neck_ and *C*_neck_: when *h*_*c*,2_ < 0, a small *r*_neck_ and large negative *C*_neck_ (i.e., sharp neck region) can result in negative *h*_*c*_. We expect to observe a preference for cell orientation along the circumferential direction (*φ* direction) for a neck region with a sharp shape (large negative *C*_neck_) and short interface length (small *r*_neck_), while a longer and smoother neck region leads to cell orientation along the *s* direction. Our simulations confirm these predictions, as shown in Fig. 3(d,e).

We next perform experiments to validate our theoretical predictions. For a bead doublet with a sharp neck and a short interfacial length (Fig. 1(f)), we observed preferential alignment of TCs along the circumferential direction (Fig. 1(h)). However, for a bead doublet with a smooth neck and a longer interfacial length, we found that TCs accumulate at the neck region, leading to a multilayer structure and bimodal orientation behavior where the inner TCs orient mainly along the circumferential direction, whereas the outer cells orient along the perpendicular direction (Fig. 3(f,g)). This bimodal distribution of cell orientation can be understood by our theory: inner cells sense a small neck radius *r*_neck_ and a sharp neck (a larger negative *C*_neck_) while outer cells sense a larger neck radius *r*_neck_ and a smoother neck (a smaller negative *C*_neck_). Thus, the formation of the multilayer structure results in variations of the effective surface curvature that TCs sense, which leads to the emergence of bimodal orientation of TCs.

To further explore the cell orientation behavior in response to the substrate curvature, we next examine a dome-like surface (Fig. 4(a,b)), which is rotationally symmetric about the *z* axis and can be characterized by a curvature profile *C*_*ss*_(*s*) = *C*_*m*_ cos(*πs/L*_*m*_) for *s < L*_*m*_ and *C*_*ss*_(*s*) = 0 for *L*_*m*_ *< s < L*_*m*_ + *L*_0_ (Fig. 4(b)). We find that increasing the dome curvature *C*_*m*_ leads to spontaneous flows, consistent with previous studies [22]. With the same curvature-sensing parameters of *h*_*c,i*_ as in simulations of the curved strip geometry and the hour-glass geometry, we observe that cells at the neck region (defined as the region where *C*_*ss*_ < 0, thus also saddle-shaped region) tend to orient along the circumferential direction (i.e., the *φ* direction) (Fig. 4(c,d)). By contrast, with a constant curvature-sensing parameter, *h*_*c*_ = const (i.e., *h*_*c*,1_ = 0 and *h*_*c*,2_ = 0 here), cells at the neck region tend to orient along the weft direction (i.e., the *s* direction). Importantly, decreasing the dome curvature *C*_*m*_ leads to cells at the neck region being more randomly orientated, sometimes along the *s* direction.

**FIG. 4.**
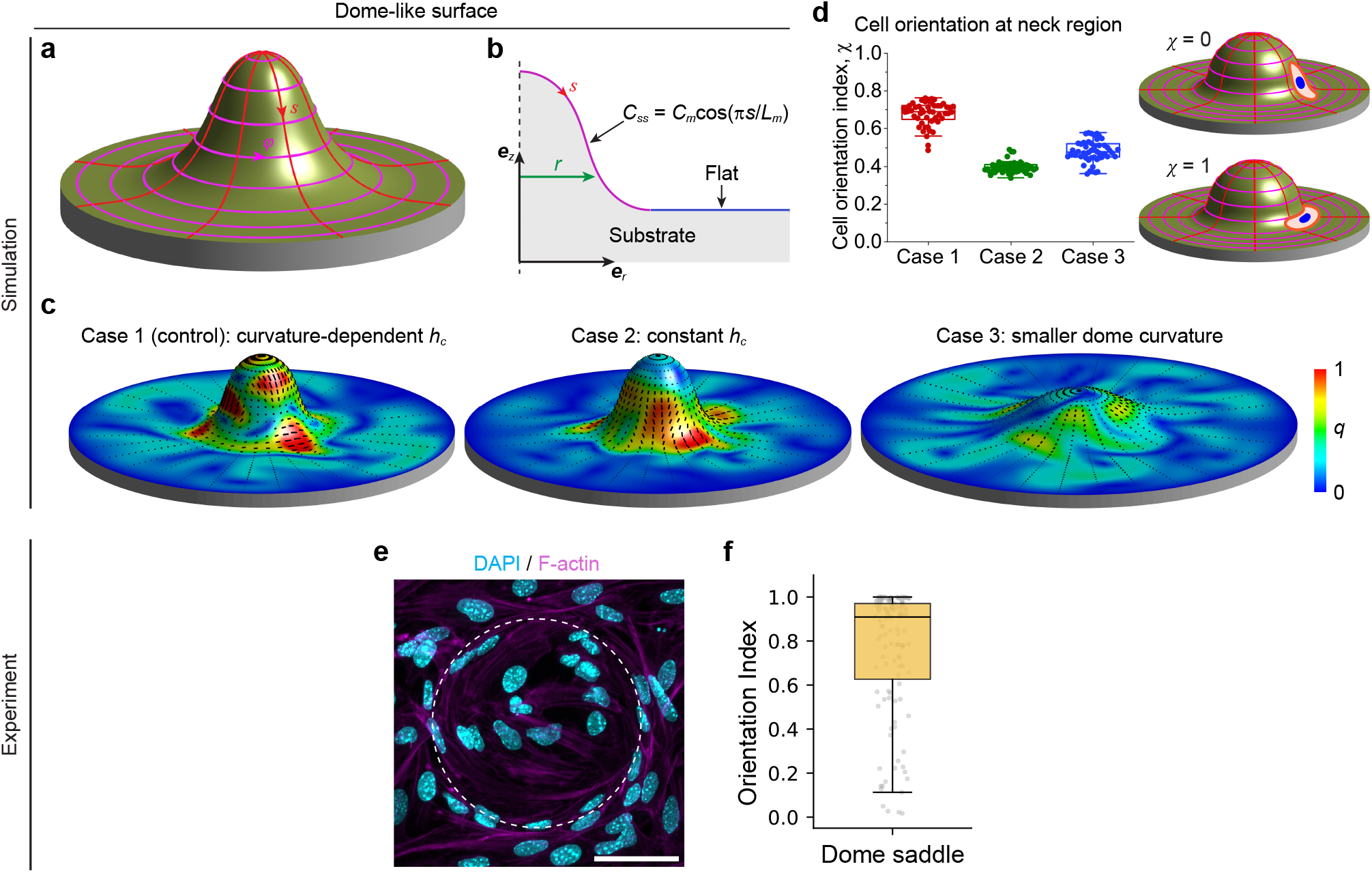
Cell orientation on a dome-like geometry: (a-d) simulation; (e-f) experiment. (a) 3D view of a dome-like surface. Here, we also show the surface coordinates (*s, φ*). (b) Cross-sectional view of the dome-like surface geometry used in simulation. It is composed of two regions, including a curved region and a flat plane of length *L*_0_. The former, curved region is characterized by its curvature profile *C*_*ss*_ = *C*_*m*_ cos(*πs/L*_*m*_) with *C*_*m*_ quantifying the curvature magnitude and *L*_*m*_ being the arc length. (c, d) Comparison of the cell orientation map (c) and the cell orientation index for cells at the neck region (d), for three cases: case 1 (control case), curvature-dependent *h*_*c*_ (see Eq. (3)); (2) case 2, constant *h*_*c*_; (3) case 3, smaller dome curvature (compared to case 1). In (c), the color code represents the magnitude of cell elongation *q*; the black lines indicate the local direction of cell orientation. (e) Maximum projection of fluorescence image of TCs on dome substrates. The white dotted circle indicates the neck region of the dome. Scale bar, 50 *µ*m. (f) Boxplot of cell orientation index at the neck region of dome substrates. *n* = 119 nuclei from 7 domes in two independent experiments were analysed.

To validate our theoretical predictions, we prepared substrates with dome-like structures (see Methods). As TCs reached confluency (approximately 72 hours after seeding), we observed that TCs at the neck region tend to orient along the circumferential direction (i.e., the *φ* direction), as shown in Fig. 4(e), in line with our simulation result with a curvature-dependent *h*_*c*_. This again suggests that a curvature-dependent *h*_*c*_ is required to account for the orientation of TCs over curved substrates.

### Phase diagrams of cell orientation regulated by substrate curvature and curvature-sensing parameters

Overall, we summarize our findings of the curvature-induced cell re-orientation response by projecting the median cell orientation index *χ* onto a phase space defined by the mean curvature *H* and the Gaussian curvature *K*, as shown in Figs. 5(a,b). It clearly demonstrates that both the mean curvature *H* and the Gaussian curvature *K* regulate cell orientation. Specifically, in saddle-shaped regions with negative Gaussian curvature, there exists a regime in which the dominant cell orientation switches as the mean curvature changes. Also, in some parameter regions, a small decrease in the Gaussian curvature *K* from zero (corresponding to a hemicylindrical surface) significantly enhances a collectively coherent cell orientation.

**FIG. 5.**
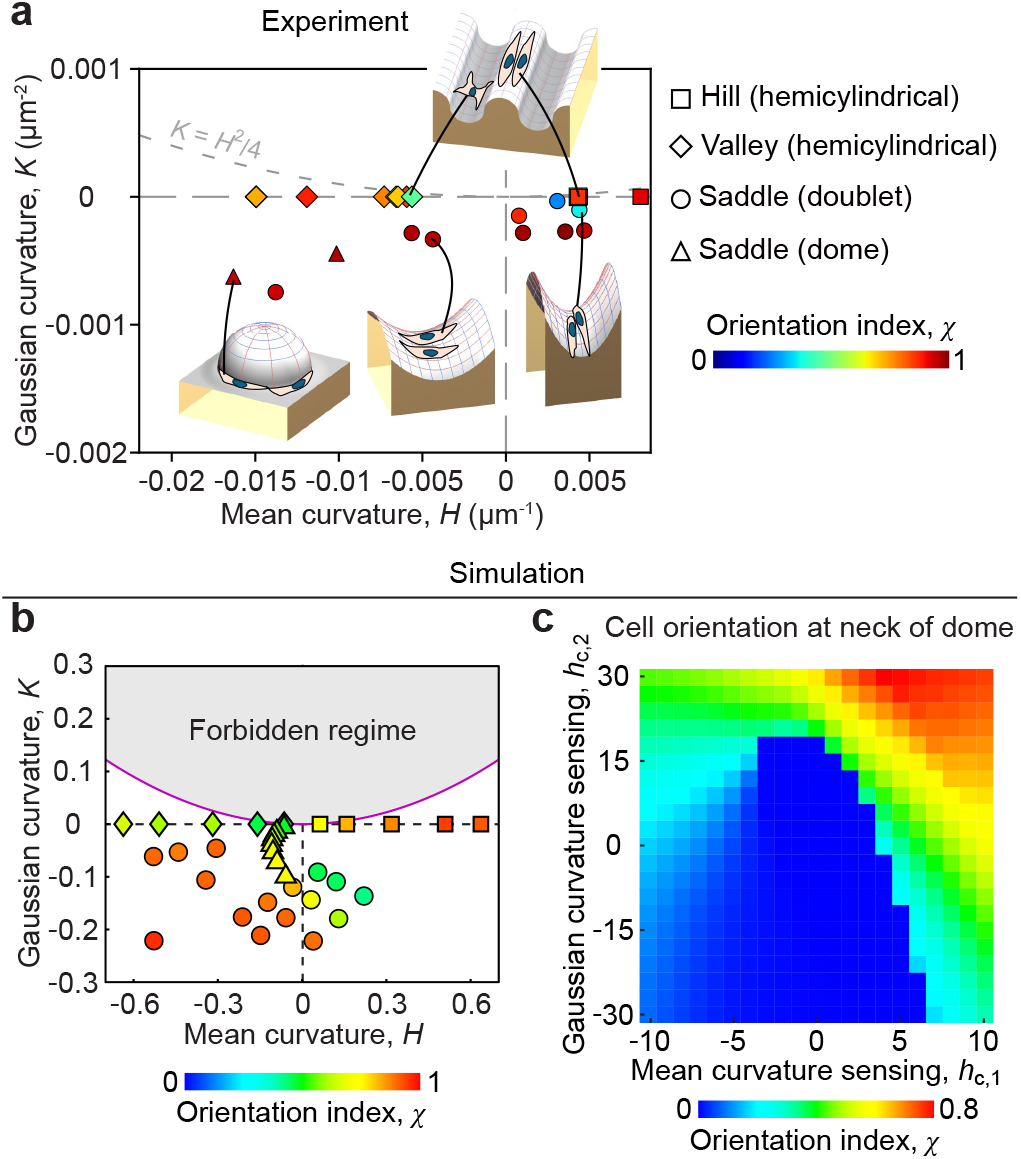
Regulation of cell orientation by distinct curvatures and curvature-sensing parameters. (a, b) Experimental data (a) and theoretical simulation (b) for cell orientation regulated by the mean curvature *H* and the Gaussian curvature *K* in various curved geometries. Symbols indicate the type of substrates used in the present study, while the heatmap shows the median of the cell orientation index extracted from each image or simulation. (c) Phase diagram of the cell orientation index at the neck region of a dome, regulated by the curvature-sensing parameters *h*_*c*,1_ and *h*_*c*,2_.

In addition, we theoretically explore how the mean curvature-sensing parameter *h*_*c*,1_ and the Gaussian curvature-sensing parameter *h*_*c*,2_ regulate cell orientation. Here, for simplicity, we focus on the neck region of a dome geometry. We represent a phase diagram of the cell orientation index *χ* in Fig. 5(c). It shows that the cell orientation is regulated by both *h*_*c*,1_ and *h*_*c*,2_. Varying *h*_*c*,1_ or *h*_*c*,2_ from zero to large values leads to a switch of cell orientation at saddle-shaped regions.

### Local cell accumulation can be explained by curvature-induced traction force

In our experiments, besides the orientation response of TCs to the substrate curvature, we also observed that TCs tend to migrate along the curvature gradient, a phenomenon known as curvotaxis [1, 40, 41]. Specifically, we observed that TCs initially placed at the valley region (negative curvature) tend to migrate toward the hill region (positive curvature) [42]. We propose that such a curvotaxis behavior of TCs can be described by a curvotaxis traction force, 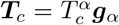. In general, 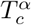 depends on 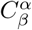. Here, inspired by our experiments, we express 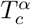 in a simple formalism:

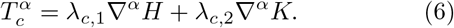

The first term in Eq. (6) accounts for the motion tendency of cells along the gradient of the mean curvature *H* (referred to as mean curvotaxis) with a strength parameter *λ*_*c*,1_ (Fig. 6(c)): *λ*_*c*,1_ > 0 for cells that tend to move from a lower mean curvature region to a higher mean curvature region, e.g., the TCs in our experiments; otherwise, *λ*_*c*,1_ < 0, e.g., MDCK cells [40]. The second term in Eq. (6) accounts for the migration tendency of cells along the gradient of the Gaussian curvature *K* (referred to as Gaussian curvotaxis) with a strength *λ*_*c*,2_.

**FIG. 6.**
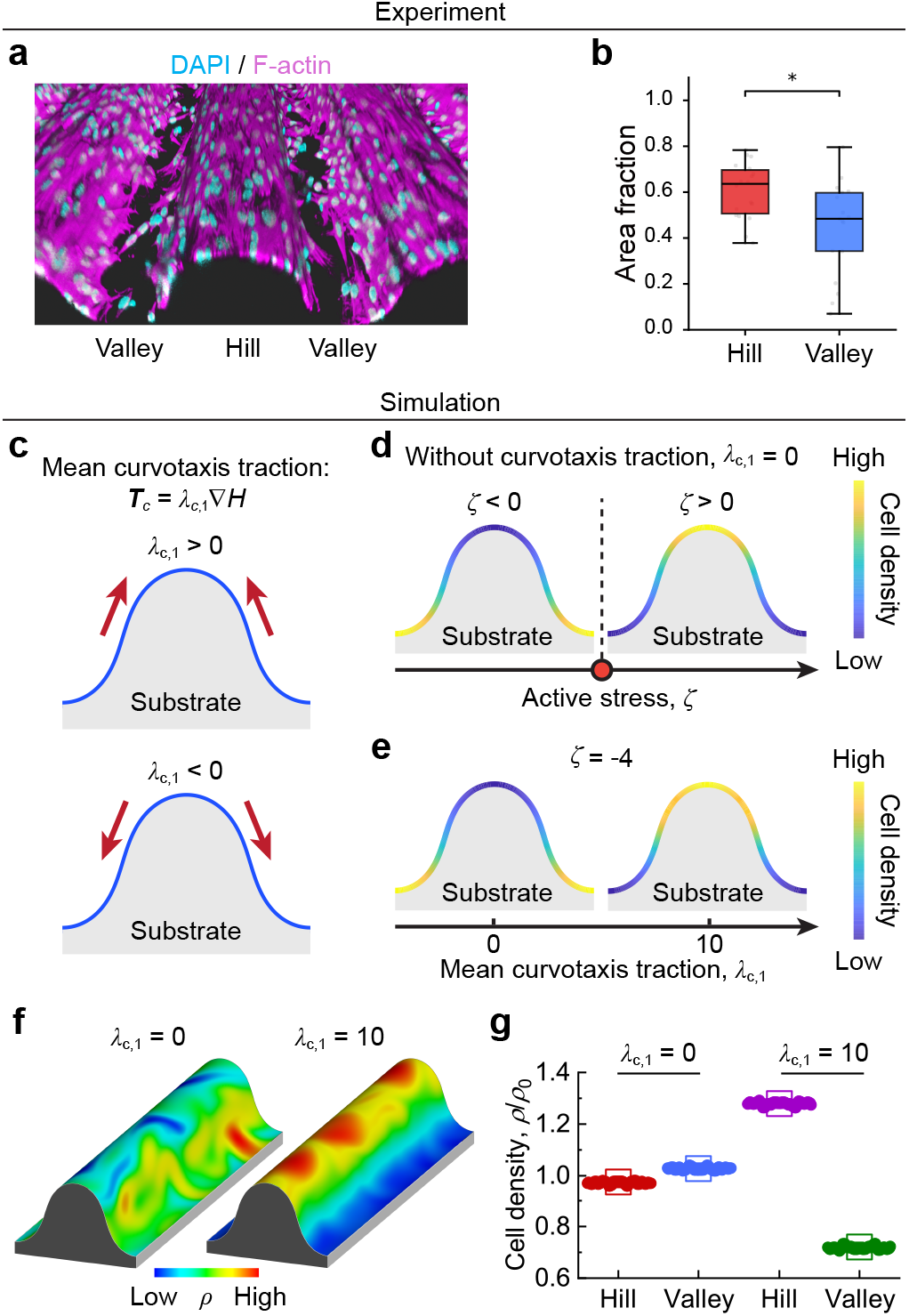
Roles of the cellular active stress *ζ* and the mean curvotaxis traction *λ*_*c*,1_ in regulating the cell density profile. 3D rendering image of TCs on 100 µm hemicylindrical substrates. Images were obtained 72 h after seeding. (b) Comparison of the cell densities at the hill region and at the valley region on 100 *µ*m hemicylindrical substrates. Statistical significance was assessed using a two-sided Mann–Whitney U test. ^∗^*P* < 0.05; *n* = 18 regions for hill, *n* = 17 regions for valley from 3 independent experiments. (c) Sketch of the mean curvotaxis traction force. (d) Schematic of the cell density profile regulated by the cellular active stress *ζ*. TCs in experiments correspond to the regime of *ζ* < 0. (e) The cell density profile regulated by the mean curvotaxis traction *λ*_*c*,1_ with *ζ* = −4. (f, g) Numerical simulation. (f) Comparison of the cell density map for two cases: (*left*) *λ*_*c*,1_ = 0; (*right*) *λ*_*c*,1_ = 10. Here, the color code represents the cell density *ρ*. (g) Comparison of the cell density at the hill region and at the valley region, corresponding to (f).

By introducing such a curvotaxis traction force (Eq. (6)) into the force balance equation (4) (Supporting Information, Sec. I), we explore how it regulates the cell density profile. We first examine the curved strip surface. For such a curved surface, we obtain an analytical expression of the cell density profile at the steady-state (see Supporting Information, Sec. IIA):

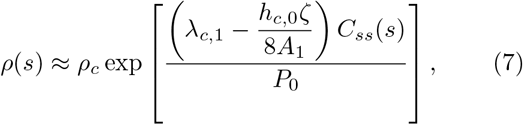

with *ρ*_*c*_ a constant, determined by the mass conservation constraint. Applying the theoretical prediction Eq. (7) to TCs (*h*_*c*,0_ > 0, *h*_*c*,1_ > 0, *ζ* < 0, *λ*_*c*,1_ > 0), we found that the cellular active stress (*ζ*) results in an accumulation of TCs at the valley region while the mean curvotaxis traction (*λ*_*c*,1_) results in accumulation of TCs at the hill region (Fig. 6(d, e)). Combined with our experimental observation that TCs accumulated at the hill region (Fig. 6(a,b)), it indicates that for TCs the mean curvotaxis traction (*λ*_*c*,1_) dominates over the cellular active stress (*ζ*). We performed numerical simulations to validate our analytical predictions. As expected, our simulations agree well with our theoretical prediction and experiments (Fig. 6(e,f)).

Equation (7) also indicates that the substrate curvature magnitude (|*C*_*ss*_|) affects the cell density profile: decreasing the curvature can decrease the cell density difference between the hill and valley regions. To experimentally validate the regulating role of substrate curvature, we cultured TCs over a hemicylindrical substrate of lower curvature (*C*_*ss*_ = 0.005 *µ*m^−1^ vs. *C*_*ss*_ = 0.01 *µ*m^−1^). In this case, we still observed accumulation of TCs at the hill region, but the cell density difference between the hill and valley regions is lower in the case of lower curvature substrates (Fig. S1), in line with our theoretical prediction.

We next performed numerical simulations to examine the cell density profiles on hourglass and dome-like surfaces (Fig. S2). In particular, we explored how the mean curvotaxis traction *λ*_*c*,1_ and the Gaussian curvotaxis traction *λ*_*c*,2_ regulate the cell density profile. In experiments, we observed TC accumulation at saddle-shaped regions of both the hourglass surface and the dome-like surface (Fig. S3 and Fig. S4). Our above study on the hemicylindrical surface already indicates that *λ*_*c*,1_ > 0 for TCs. Here, we further find that only with a negative value of the Gaussian curvotaxis traction (i.e., *λ*_*c*,2_ < 0) can we numerically reproduce cell accumulation at saddle-shaped regions (Fig. S2). This further augments the critical role of Gaussian curvature in driving TC flow.

Notably, our theory predicts distinct flow patterns of cell layers over curved surfaces. For example, on a hemi-cylindrical surface (Fig. 1(a)), we obtain a transition of flow patterns from a non-flowing state to a shear flow state, when the cell activity *ζ* or the surface curvature *C*_0_ exceeds a critical value (Fig. S5). This agrees with previous studies [22, 35, 43]. We further find that the curvature-sensing parameters (*h*_*c*,0_, *h*_*c*,1_, *h*_*c*,2_) regulate the flow patterns (Fig. S5 and Fig. S6).

## DISCUSSION AND CONCLUSION

In this work, we presented a hydrodynamic theory of active nematic gel on curved surfaces to describe the collective cell alignment and density variations of ovarian stromal-like TCs on various curved geometries. Specifically, we showed that a generalised curvature-sensing coefficient containing non-linear curvature-dependent parameters is sufficient to recapitulate TC orientation on hemicylindrical, hourglass- and dome-like substrates with various combinations of the mean curvature and the Gaussian curvature. By introducing a curvotaxis traction force, our model predicts an accumulation of TCs on convex surfaces of hemicylindrical substrates and saddle-shaped regions on bead doublet surfaces and dome-like surfaces, consistent with experimental observations.

There have been many studies investigating how cells migrate and align on hemicylindrical surfaces (with zero Gaussian curvature). For example, T cells have been reported to migrate along the concave surfaces of sinusoidal wavy substrates [44]. By contrast, Werner *et al*. [9, 45] showed that bone marrow stromal cells align and migrate along the cylinder axis with zero curvature, consistent with our observation of TC alignment on hemicylinder substrates. Similarly, fibroblasts and vascular smooth muscle cell monolayer displayed preferential alignment along the axial direction on cylinders [46], suggesting that the behavior is not confined to specific types of stromal cells. Other studies on convex and concave surfaces that have positive Gaussian curvatures showed that mesenchymal stem cells are more migratory on concave surfaces than on convex surfaces, with the latter inducing osteogenic differentiation due to greater nuclear deformation [47]. Interestingly, cells on cell-size concave/convex microwells exhibit an opposite response [40], where they tend to avoid convex hills and remain largely in concave valleys, suggesting that the relative size of cells to the substrate curvature is an important consideration in understanding tissue alignment and migration.

By contrast, how cells behave on surfaces with negative Gaussian curvature is less well understood. A previous study showed that fibroblasts on domes can orient themselves and migrate along the circumferential direction [48], similar to our observation of TCs on domes and bead doublets. In the case of bead doublets, we observed that with increasing number of TCs lying at the neck in a circumferential manner, the outermost layer of TCs experienced a decrease in local curvature. Effectively, these cells sense a more ‘cylinder-like’ geometry, thereby realigning along the zero curvature axis, that is, across the neck (see Fig. 3g). These studies provide strong evidence that changes in the surface topology can elicit distinct and novel transitions in tissue nematic order and flow.

Past studies have suggested that cell alignment in the zero curvature axis of a cylindrical surface is likely attributed to the minimization of bending energy in the circumferential direction. Deviation from this direction forces the cell to adapt to a more bent configuration, which is energetically unfavorable. However, it remains unclear why cells would prefer to align along the circumferential direction in the saddle-shaped regions, although there is some indication that the different polarisation of apical versus basal stress fibers for cells at the saddle-shaped region may minimize the free energy [48]. Future studies on molecular staining of stress fibers and acto-myosin contractility will likely shed light on this aspect.

Another open question pertains to the active migration of TCs onto the hills. While we have shown that a curvature-induced active cell traction is essential to drive this behavior, the molecular and cellular mechanisms driving this process remain unknown. Such curvotaxis will likely involve focal adhesion dynamics, nuclear shape, and actin cytoskeleton remodelling [49]. Another missing ingredient in our model is the role of extra-cellular matrix (ECM) remodelling. Mesenchymal cells, such as fibroblasts or myoblasts, are known to continually secrete ECM to maintain tissue structural integrity and homeostasis. We have recently shown that functional TC contractility directly contributes to the synthesis of fibronectin and hyaluronic acid in TC layers around ovarian follicles [34, 42]. Future refinement of the model to incorporate additional cell-substrate adhesion terms and possible crosstalk between cells and ECM [50, 51], combined with experimental validations, will shed light on how the crosstalk between curvature, ECM, and active cell traction leads to robust tissue patterning and flow. We note that in describing the evolution of cell density, our model has ignored the effects of cell division or apoptosis. However, experimentally, we did observe more cell divisions at the hills compared to those at valleys [42], suggesting that cell accumulation could be a combined effect of active curvotaxis and increased cell division at the hill. Future theoretical work may incorporate non-conservative elements such as cell density changes due to cell proliferation and death.

Despite these limitations, our theoretical framework provides a new approach to understanding active cell migration and the emergent nematic order on substrates of various curvatures. The generic nature of our model also implies that it can, in principle, be applied to study tissue dynamics of any contractile cells with shape anisotropy, which needs to be validated by future experiments.

## MATERIALS AND METHODS

### Animals

Mice were group housed in individually ventilated cages with access to water and food under a 12-hr light/12-hr dark cycle. Mouse rooms were maintained at 18 −25 °C and 30 −70% relative humidity. C57BL/6NTac female mice, aged P25 – P28, were euthanized by carbon dioxide asphyxiation followed by cervical dislocation. All animals were bought from Singapore InVivos. Ovaries were then dissected from the mice and transferred to an isolation buffer consisting of Leibovitz’s L15 medium (Thermo Fisher, 21083027) supplemented with 3 mg/ml Bovine Serum Albumin (BSA, Sigma, A9647). All mice care and use were approved by the Institutional Animal Care and Use Committee (IACUC) at the National University of Singapore.

### Primary TC isolation and culture

The assay to isolate primary TCs has been established in our lab [34], following adaptation of a previous study [52]. In brief, freshly isolated ovaries were washed three times in isolation buffer at room temperature and poked with a needle to burst the follicles and release the granulosa cells under the stereomicroscope. The ovaries were punctured until intact follicular architecture was not visible. The buffer with the granulosa cells was centrifuged at 94× g for 5 minutes. The pellet was washed twice and resuspended in McCoy’s 5A medium (16600082, Gibco) supplemented with 5% FBS and 1% Penicillin-Streptomycin. A digestion buffer comprising 0.05 mg*/*mL activated DNase I (D4263, Sigma) and 10 mg*/*mL collagenase IV (17104-019, Thermo Fisher) in M199 (11150067, Gibco) medium was freshly made. DNase was activated by mixing DNase I with HBSS (14025-092, Gibco) in a 1 : 1 ratio. After mechanical disruption of the follicles to release the intrafollicular GCs, the remaining tissue was washed twice with isolation buffer and transferred to the digestion buffer (200 *µ*L of buffer was used per ovary). The tissue was incubated at 37 °C for an hour with gentle mixing every 15 minutes using a pipette. Once completely dissolved, the solution was centrifuged at 94 × g for 5 minutes. The pellet was washed and resuspended in McCoy’s medium supplemented with 5% FBS and 1% Pen-Strep to obtain the theca/stroma cells. The cells were then seeded onto 6-well cell culture dishes (ThermoFisher Scientific, NunclonTM Delta Surface) and incubated at 37 °C, 5% CO2, and 95% humidity overnight before further experiments. The purity of TCs isolated through this method has been demonstrated in our previous study [34].

### Follicle isolation and culture

Follicles were mechanically isolated from dissected ovaries under the stereomicroscope attached to a thermal plate using tweezers in isolation buffer at 37 °C. Growth medium, consisting of MEM*α* GlutaMAX (Thermo Fisher, 32561037) supplemented with 10% Fetal Bovine Serum (FBS, Thermo Fisher, 10082147), 1% Penicillin-Streptomycin (Thermo Fisher, 10378016), 1× Insulin-Transferrin-Selenium (ThermoFisher, 41400045), and 50 mIU/mL follicle-stimulating hormone (Sigma, F4021) was prepared. Individual follicles were transferred to a custom-made 1% agarose well mounted on a 35 mm petri dish (ThermoFisher) containing the above-mentioned growth medium. Follicles were cultured for 24 hours at 37 °C, 5% CO2, and 95% humidity until re-seeded for assays.

### TC culture on hemicylindrical and dome substrates

Custom polyacrylamide (PA) hydrogel substrates were prepared by replicating microfabricated molds with defined surface geometries. Two types of microtopographies were used: (1) hemicylindrical wave structures with half-periods of 100 and 200 *µ*m, following the design described by Ref. [53], and (2) dome-shaped structures fabricated by the MBI Microfabrication Core Facility (height ≈50 *µ*m, width ≈106 *µ*m). PA hydrogels were cast using pre-optimized acrylamide and bis-acrylamide combinations to achieve the desired stiffness. Coverslips were UV-cleaned and treated sequentially with 0.3% acetic acid and 0.5% 3-(trimethoxysilyl)propyl methacrylate (TMSPMA) in ethanol, rinsed twice in 100% ethanol, and air-dried. Immediately before casting, ammonium persulfate (APS) and tetramethylethylenediamine (TEMED) were added to the PA precursor. The mixture was dropped onto a dichlorodimethylsilane (DCDMS)–treated mould and covered with the treated coverslip to form the gel. After polymerization, the hydrogel was peeled off, washed three times with 50 mM HEPES, and incubated at room temperature for 30 min. Sulfosuccinimidyl 6-(4-azido-2-nitrophenylamino) hexanoate (Sulfo-SANPAH in 50 mM HEPES buffer) was activated under UV (UV-KUB 9, 6% power, 4 min) to enable protein coupling. The gels were rinsed three times in HEPES buffer, followed by incubation in a solution containing collagen I (bovine tail, Gibco A1064401) adjusted to 50mg*/*mL in 20 mM acetic acid for ≥3 h at room temperature or overnight at 4 °C. Collagen-coated gels were rinsed with PBS and sterilized under UV light before cell seeding. Isolated TCs (∼2.5 × 10^5^ cells per 22 × 22 mm coverslip) were seeded and cultured for 72 h before fixation with 4% paraformaldehyde.

### TC culture on hydrogel bead doublets

Gelatin methacryloyl (GelMA) hydrogel beads were fabricated using a microfluidic droplet generator. Flow rates of the continuous and dispersed phases were tuned to produce spherical beads with diameters ranging from 100 *µm* to 400 *µm*. Isolated ovarian follicles were selected to match the bead size and co-cultured in direct contact with the GelMA beads for 2–3 days, allowing TCs to encapsulate and spread the bead surface. After this precoating phase, the follicles were mechanically removed under a stereomicroscope, leaving a uniform TC layer on the bead. A newly prepared, size-matched clean bead was gently placed adjacent to the cell-coated bead, and the pair was cultured for an additional 2–3 days. During this period, spontaneous fusion occurred, forming hydrogel bead doublets fully enveloped by TCs. The samples were subsequently fixed in 4% paraformaldehyde for subsequent imaging and analysis.

### Immunofluorescence staining and imaging

4% paraformaldehyde was used to fix samples, followed by washing in buffer (1% BSA in PBS) for each experiment. TCs cultured on hemicylindrical and dome substrates were fixed after 72 hours of culture, when they reached confluency. TCs cultured on doublet substrates were fixed once they fully covered the surface, approximately 3 days after the introduction of the second bead, corresponding to 6–7 days after the start of culture. The samples were incubated in blocking–permeabilization buffer containing 3% Bovine Serum Albumin (BSA) and 0.03% Triton X-100 (Sigma, X100) at room temperature for 2–4 h. Samples were washed five times with washing buffer and stained with DAPI (2 *µ*g*/*mL, Sigma, D9542) and Alexa Fluor 633-conjugated phalloidin (1:300, Invitrogen, A22284) diluted in washing buffer for 4 h at room temperature. After staining, the samples were washed three times with washing buffer before mounting. For the dome and hemicylindrical substrates, coverslips were mounted with the cell-adhesive side facing upward. In contrast, bead doublet samples were mounted in Slow-Fade Gold Antifade Mountant (Thermo Fisher, S36940). Confocal imaging was performed using a Nikon A1R confocal laser-scanning microscope equipped with a CFI Apo LWD Lambda S 40× water-immersion objective (NA = 1.15). Z-stack images were acquired at 1–4 *µ*m intervals under identical imaging parameters across all samples.

### Curvature analysis

Surface curvature was quantified by extracting orthogonal cross-sections from 3D z-stack images in Fiji. Circular fitting was applied to both the concave (saddle) and convex (ring or dome-like) regions to obtain the two principal curvature radii (*r*_1_, *r*_2_). The corresponding principal curvatures (*κ*_1_, *κ*_2_) were calculated as *κ*_1_ = 1*/r*_1_ and *κ*_2_ = 1*/r*_2_, with the sign convention that concave surfaces are negative and convex surfaces positive. Mean curvature (*H*) and Gaussian curvature (*K*) were computed as

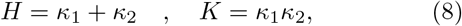

respectively. Data from all samples were aggregated and visualized using a custom Python-based pipeline that grouped results by substrate type (saddle doublet, saddle dome, hill, valley) and generated *H*–*K* phase diagrams to classify local surface geometries.

### Nuclear orientation and density analysis

Analysis of nuclear orientation and density on the hemicylinder was analyzed separately for the hill and valley regions. Nuclei within each respective region were projected using a Z-intensity projection, and their orientation was analyzed using the Analyze particles function in ImageJ. Because TCs did not reach confluency, particularly in the valley region, we defined the area fraction based on the coverage of the actin signal over the respective surface areas of the hill and valley. As the actin signal intensity was heterogeneous between the hill and valley regions, contrast adjustment was performed prior to generating binary masks. The surface areas of the hill and valley were then calculated using the measured diameter and length of the hemicylindrical substrate. Nuclear orientation and density on dome and bead doublet substrates were performed using a custom-built pipeline based on Python and the skimage, scipy, and numpy libraries. First, 3D binary masks for nuclei and actin were generated from 3D images (DAPI and actin channels) by sequentially applying Otsu’s thresholding, morphological opening with a physical spherical structuring element, small object removal, and binary hole filling.

For dome and bead doublets, reference sphere centers were estimated by spherical fitting to actin-derived surface point clouds. In doublets, actin surface points were further separated into two clusters to estimate the two sphere centers. The 3D long axis of each nucleus was determined by applying Principal Component Analysis (PCA) to its segmented physical coordinate cloud. Specifically, Singular Value Decomposition (SVD) was performed on the coordinates, and the first principal component vector, corresponding to the direction of maximum variance, was defined as the nuclear orientation vector.

The cell orientation index *χ* was defined consistently across the doublet, cylinder, and dome geometries based on the local coordinate system. Specifically, the orientation index *χ* was calculated as the absolute sine of local cell orientation angle *θ*, i.e., *χ* = |sin *θ* |, so that it ranges from 0 to 1. In each geometry, the angle *θ* was measured between the nuclear long axis and the local meridional direction. For example, for a hemicylindrical surface, *θ* is the angle between the orientation of a cell nucleus and the curvature direction of the hemicylindrical substrate.

For dome substrates, nuclear density was quantified using an area-normalized approach. Specifically, the number of nuclei within each region was divided by the corresponding surface area, which was computed from a reconstructed 3D mesh of the substrate derived from the actin signal. Regions were defined based on height relative to a fitted reference plane as dome (>20 *µ*m), saddle (5–20 *µ*m), and flat (<5 *µ*m). For bead-doublets, density was likewise quantified using an area-normalized approach. The number of nuclei within each region was divided by the corresponding mesh-derived surface area, calculated separately for the neck and surface regions within the bottom half of the substrate, where signal attenuation was minimal, and segmentation could be performed reliably.

### Model and simulation

Details of the model and simulation schemes over various curved substrate surfaces are presented in the Supporting Information.

### Statistical analysis

Unless otherwise stated, all statistical analyses were performed using Python 3. Data were analyzed using the two-tailed Mann-Whitney U test to determine statistical significance between groups. All experiments were conducted with at least two independent biological replicates (animal replicates), and each experiment included at least five technical replicates to ensure reproducibility.

## Author contributions

S.Z.L. and C.J.C. designed research; S.Z.L., K.T., B.H.N., C.H.L., J.S.W., and C.J.C. performed research; K.T. and S.Z.L. analyzed data; Z.Z. and E.L.S.E. contributed the gelatin beads; S.Z. L., K.T., and C.J.C. wrote the paper.

## Acknowledgement

S.Z.L. acknowledges support from the National Natural Science Foundation of China (Grant No. 12502367), Fundamental Research Funds for the Central Universities, Sun Yat-sen University (Grant No. 25hytd014), Research Center for Magnetoelectric Physics of Guangdong Province (Grants 2024B0303390001), and Guangdong Provincial Key Laboratory of Magnetoelectric Physics and Devices (Grants 2022B1212010008). C.J.C. acknowledges support from the Ministry of Education under the Research Centres of Excellence programme through the Mechanobiology Institute and the Department of Biological Sciences at the National University of Singapore, the Ministry of Education Tier2 grant (T2EP30222-0026), National Research Foundation Mid-size Grant (NRF-MSG-2023-0001), and the Bia-Echo Asia Centre for Reproductive Longevity and Equality (ACRLE) at the National University of Singapore. C.J.C. acknowledges the support of the Singaporean Teaching and Academic Research Talent Inauguration Grant (START). We thank Cheng-Kuang Huang for providing the hemicylindrical moulds, and Yuting Lou for initial discussion of data. We thank the support of the Singapore Microscopy and Bioimaging Analysis (SiMBA, MBI) core for microscopy, and the MBI Microfabrication core.

