## Supplementary information for "Dual curvature sensing governs cell orientation and curvotaxis"

### Supporting Information

Here, we provide details of the model and simulation. In Sec. I, we give the governing equations for a compressible, active nematic layer adhering to a general curved substrate surface. In Sec. II, we give the detailed governing equations and simulation schemes for some typical curved substrate surfaces explored in the present study, including a hemicylindrical surface, an hourglass surface, and a dome-like surface. In Sec. III, we give some supplementary figures associated with the main text.

#### CONTENTS


I. Governing equations for general curved substrate surfaces 1
  A. Substrate surface geometry 1
  B. Governing equations 2

II. Some typical curved substrate surfaces studied 4
  A. Hemicylindrical geometry 4
    1. Governing equations 4
    2. Numerical simulation 5
    3. Theoretical analysis 7
  B. Hourglass geometry 8
    1. Governing equations 9
    2. Numerical simulation 10
  C. Dome-like geometry 11
    1. Governing equations 12
    2. Numerical simulation 13

  References 13

III. Supplementary figures for the main text 14


#### I. GOVERNING EQUATIONS FOR GENERAL CURVED SUBSTRATE SURFACES

##### A. Substrate surface geometry

Consider a generic curved surface embedded in $\mathbb{R}^3$. Let $\boldsymbol{r} = \boldsymbol{r}(x^1, x^2)$ be the position on such a generic surface parametrized by the coordinates $(x^1, x^2)$. Let

$$\boldsymbol{g}_\alpha = \frac{\partial \boldsymbol{r}}{\partial x^\alpha}, \tag{S1}$$

be a basis of covariant vectors on the tangent plane. Then, the surface metric tensor reads,

$$\boldsymbol{g} = g_{\alpha\beta}\boldsymbol{g}^\alpha \otimes \boldsymbol{g}^\beta = g^{\alpha\beta}\boldsymbol{g}_\alpha \otimes \boldsymbol{g}_\beta = \boldsymbol{g}^\alpha \otimes \boldsymbol{g}_\alpha = \boldsymbol{g}_\alpha \otimes \boldsymbol{g}^\alpha, \tag{S2}$$

where $g_{\alpha\beta} = \boldsymbol{g}_\alpha \cdot \boldsymbol{g}_\beta$ and $g^{\alpha\beta} = \boldsymbol{g}^\alpha \cdot \boldsymbol{g}^\beta$ with $\boldsymbol{g}^\alpha$ being a basis of contravariant vectors on the tangent plane, satisfying $\boldsymbol{g}^\alpha \cdot \boldsymbol{g}_\beta = \delta^\alpha_\beta$. $g^{\alpha\beta}$ and $g_{\alpha\beta}$ satisfy, $(g^{\alpha\beta})(g_{\alpha\beta}) = I_2$ with $I_2$ being the second-order identity matrix. Then, $\boldsymbol{g}^\alpha$ can be calculated by $\boldsymbol{g}^\alpha = g^{\alpha\beta}\boldsymbol{g}_\beta$.

The extrinsic curvature tensor reads,

$$\boldsymbol{C} = C_{\alpha\beta}\boldsymbol{g}^\alpha \otimes \boldsymbol{g}^\beta = C^{\alpha\beta}\boldsymbol{g}_\alpha \otimes \boldsymbol{g}_\beta = C^\alpha_\beta \boldsymbol{g}_\alpha \otimes \boldsymbol{g}^\beta, \tag{S3}$$

where

$$C_{\alpha\beta} = -\frac{\partial^2 \boldsymbol{r}}{\partial x^\alpha \partial x^\beta} \cdot \boldsymbol{m}, \tag{S4}$$

with $\boldsymbol{m} = (\boldsymbol{g}_1 \times \boldsymbol{g}_2)/|\boldsymbol{g}_1 \times \boldsymbol{g}_2|$ being the normal vector of the curved surface. Then, the mean curvature $H$ and the Gaussian curvature $K$ can be calculated by,

$$H = \mathrm{tr}(C^\alpha_\beta) = C^\alpha_\alpha, \tag{S5}$$

and

$$K = \det(C^\alpha_\beta), \tag{S6}$$

where $C^\alpha_\beta = g^{\alpha\gamma} C_{\gamma\beta}$.

In addition, the Christoffel symbols (denoted $\Gamma^\gamma_{\alpha\beta}$) are related to the gradient of the surface metric tensor, defined as:

$$\Gamma^\gamma_{\alpha\beta} = \frac{1}{2} g^{\gamma\lambda} \left( \frac{\partial g_{\alpha\lambda}}{\partial x^\beta} + \frac{\partial g_{\beta\lambda}}{\partial x^\alpha} - \frac{\partial g_{\alpha\beta}}{\partial x^\lambda} \right). \tag{S7}$$

##### B. Governing equations

In the section, we give the detailed governing equations for the nematic tensor field $\boldsymbol{Q} = Q^\alpha_\beta \boldsymbol{g}_\alpha \otimes \boldsymbol{g}^\beta$, the flow field $\boldsymbol{v} = v^\alpha \boldsymbol{g}_\alpha$, and the cell density field $\rho$.

*Nematic tensor dynamics.* – The dynamic equation of the nematic tensor $Q^\alpha_\beta$ reads:

$$\frac{\partial Q^\alpha_\beta}{\partial t} = -v^\gamma \nabla_\gamma Q^\alpha_\beta - Q^\gamma_\beta \Omega^\alpha_\gamma + \Omega^\gamma_\beta Q^\alpha_\gamma + \lambda E^\alpha_\beta + \Gamma H^\alpha_\beta. \tag{S8}$$

Here, $\nabla_\gamma$ is the covariant derivative; specifically, $\nabla_\gamma Q^\alpha_\beta = \partial Q^\alpha_\beta / \partial x^\gamma + Q^\omega_\beta \Gamma^\alpha_{\gamma\omega} - Q^\alpha_\omega \Gamma^\omega_{\gamma\beta}$, with $\Gamma^\gamma_{\alpha\beta}$ being the Christoffel symbols, given by Eq. (S7). Using the Christoffel symbols, we can derive the detailed expressions of the strain rate tensor $E^\alpha_\beta = (\nabla_\beta v^\alpha + \nabla^\alpha v_\beta - \nabla_\gamma v^\gamma g^\alpha_\beta)/2$ and the vorticity tensor $\Omega^\alpha_\beta = (\nabla^\alpha v_\beta - \nabla_\beta v^\alpha)/2$ as below:

$$E^\alpha_\beta = \frac{1}{2} \left[ \frac{\partial v^\alpha}{\partial x^\beta} + \Gamma^\alpha_{\beta\omega} v^\omega + g^{\alpha\gamma} \frac{\partial g_{\beta\lambda}}{\partial x^\gamma} v^\lambda + g^{\alpha\gamma} g_{\beta\lambda} \frac{\partial v^\lambda}{\partial x^\gamma} - g^{\alpha\gamma} g_{\omega\lambda} \Gamma^\omega_{\gamma\beta} v^\lambda - g^\alpha_\beta \frac{\partial v^\gamma}{\partial x^\gamma} - g^\alpha_\beta \Gamma^\gamma_{\gamma\omega} v^\omega \right], \tag{S9}$$

$$\Omega^\alpha_\beta = \frac{1}{2} \left( \nabla^\alpha v_\beta - \nabla_\beta v^\alpha \right) = \frac{1}{2} \left( g^{\alpha\gamma} \frac{\partial g_{\beta\lambda}}{\partial x^\gamma} v^\lambda + g^{\alpha\gamma} g_{\beta\lambda} \frac{\partial v^\lambda}{\partial x^\gamma} - g^{\alpha\gamma} g_{\omega\lambda} \Gamma^\omega_{\gamma\beta} v^\lambda - \frac{\partial v^\alpha}{\partial x^\beta} - \Gamma^\alpha_{\beta\omega} v^\omega \right). \tag{S10}$$

The total effective free energy of the nematic cell layer reads, $F = F_0 + F_C$, where

$$F_0 = \int \mathrm{d}S \left[ \frac{1}{2} A_1 Q^\alpha_\beta Q^\beta_\alpha + \frac{1}{4} A_2 \left( Q^\alpha_\beta Q^\beta_\alpha \right)^2 + \frac{1}{2} K_n \left( \nabla_\gamma Q^\alpha_\beta \right) \left( \nabla^\gamma Q^\beta_\alpha \right) \right], \tag{S11}$$

and

$$F_C = \int \mathrm{d}S \left( \frac{1}{2} h_c C^\alpha_\beta Q^\beta_\alpha \right). \tag{S12}$$

Since the nematic tensor $Q^\alpha_\beta$ is traceless (i.e., $Q^\gamma_\gamma = 0$), the curvature-sensing free energy $F_C$ can be equivalently expressed as:

$$F_C = \int \mathrm{d}S \left[ \frac{1}{2} h_c \left( C^\alpha_\beta - \frac{1}{2} C^\gamma_\gamma g^\alpha_\beta \right) Q^\beta_\alpha \right] = \int \mathrm{d}S \left( \frac{1}{2} h_c \tilde{C}^\alpha_\beta Q^\beta_\alpha \right), \tag{S13}$$

with $\tilde{C}^\alpha_\beta = C^\alpha_\beta - C^\gamma_\gamma g^\alpha_\beta / 2$ being the traceless part of the extrinsic curvature tensor $C^\alpha_\beta$. Therefore, the total effective free energy can be written as:

$$F = \int \mathrm{d}S \left[ \frac{1}{2} A_1 Q^\alpha_\beta Q^\beta_\alpha + \frac{1}{4} A_2 \left( Q^\alpha_\beta Q^\beta_\alpha \right)^2 + \frac{1}{2} K_n \left( \nabla_\gamma Q^\alpha_\beta \right) \left( \nabla^\gamma Q^\beta_\alpha \right) + \frac{1}{2} h_c \left( C^\alpha_\beta - \frac{1}{2} C^\gamma_\gamma g^\alpha_\beta \right) Q^\beta_\alpha \right]. \tag{S14}$$

Based on the above effective free energy functional, we can derive the molecular field tensor $H^{\alpha}_{\beta} = -\delta F/\delta Q^{\beta}_{\alpha}$ as:

$$\begin{aligned} H^{\alpha}_{\beta} =& - A_1 Q^{\alpha}_{\beta} - A_2 \left(Q^{\gamma}_{\lambda} Q^{\lambda}_{\gamma}\right) Q^{\alpha}_{\beta} - \frac{1}{2} h_c \left(C^{\alpha}_{\beta} - \frac{1}{2} C^{\gamma}_{\gamma} g^{\alpha}_{\beta}\right) \\ &+ K_n \left[ \begin{aligned} &\frac{1}{\sqrt{g}} \frac{\partial}{\partial x^{\lambda}} \left(\sqrt{g} g^{\lambda\gamma} \frac{\partial Q^{\alpha}_{\beta}}{\partial x^{\gamma}}\right) - \frac{1}{\sqrt{g}} \frac{\partial}{\partial x^{\gamma}} \left(\sqrt{g} g^{\gamma\lambda} \Gamma^{\chi}_{\lambda\beta} Q^{\alpha}_{\chi}\right) + \frac{1}{\sqrt{g}} \frac{\partial}{\partial x^{\gamma}} \left(\sqrt{g} g^{\gamma\lambda} \Gamma^{\alpha}_{\lambda\chi} Q^{\chi}_{\beta}\right) \\ &- g^{\gamma\lambda} \Gamma^{\chi}_{\lambda\beta} \frac{\partial Q^{\alpha}_{\chi}}{\partial x^{\gamma}} + g^{\gamma\lambda} \Gamma^{\alpha}_{\lambda\chi} \frac{\partial Q^{\chi}_{\beta}}{\partial x^{\gamma}} - 2 g^{\gamma\lambda} \Gamma^{\alpha}_{\gamma\omega} \Gamma^{\chi}_{\lambda\beta} Q^{\omega}_{\chi} + g^{\gamma\lambda} \Gamma^{\chi}_{\gamma\omega} \Gamma^{\alpha}_{\lambda\chi} Q^{\omega}_{\beta} + g^{\gamma\lambda} \Gamma^{\omega}_{\gamma\beta} \Gamma^{\chi}_{\lambda\omega} Q^{\alpha}_{\chi} \end{aligned} \right]. \end{aligned} \tag{S15}$$

Using the Christoffel symbols, the dynamic equation of the nematic tensor $Q^{\alpha}_{\beta}$ (Eq. (S8)) can be further expressed as:

$$\frac{\partial Q^{\alpha}_{\beta}}{\partial t} = -\left(v^{\gamma} \frac{\partial Q^{\alpha}_{\beta}}{\partial x^{\gamma}} + \Gamma^{\alpha}_{\gamma\omega} v^{\gamma} Q^{\omega}_{\beta} - \Gamma^{\omega}_{\gamma\beta} v^{\gamma} Q^{\alpha}_{\omega}\right) - Q^{\gamma}_{\beta} \Omega^{\alpha}_{\gamma} + \Omega^{\gamma}_{\beta} Q^{\alpha}_{\gamma} + \lambda E^{\alpha}_{\beta} + \Gamma H^{\alpha}_{\beta}, \tag{S16}$$

where $E^{\alpha}_{\beta}$, $\Omega^{\alpha}_{\beta}$, and $H^{\alpha}_{\beta}$ are given by Eqs. (S9), (S10), and (S15), in turn.

*Force balance equation.* – Considering the stress gradient, the substrate friction, and the curvotaxis traction force, the force balance equation reads:

$$\nabla^{\beta} \sigma^{\alpha}_{\beta} - \xi v^{\alpha} + \lambda_{c,1} \nabla^{\alpha} H + \lambda_{c,2} \nabla^{\alpha} K = 0, \tag{S17}$$

where the stress tensor,

$$\sigma^{\alpha}_{\beta} = -P g^{\alpha}_{\beta} + 2\eta E^{\alpha}_{\beta} - \lambda H^{\alpha}_{\beta} + Q^{\alpha}_{\gamma} H^{\gamma}_{\beta} - H^{\alpha}_{\gamma} Q^{\gamma}_{\beta} - \zeta Q^{\alpha}_{\beta}. \tag{S18}$$

Note that in our present study, we first focus on the cell orientation profile and do not consider the curvotaxis traction force; we next examine the cell accumulation behavior and include the curvotaxis traction force in our model. We find that including the curvotaxis traction force does not significantly affect the cell orientation behaviors in response to the substrate surface curvature. Using the Christoffel symbols, the force balance equation Eq. (S17) can be further expressed as,

$$g^{\beta\gamma} \frac{\partial \sigma^{\alpha}_{\beta}}{\partial x^{\gamma}} + g^{\beta\gamma} \Gamma^{\alpha}_{\gamma\omega} \sigma^{\omega}_{\beta} - g^{\beta\gamma} \Gamma^{\omega}_{\gamma\beta} \sigma^{\alpha}_{\omega} - \xi v^{\alpha} + \lambda_{c,1} g^{\alpha\beta} \frac{\partial H}{\partial x^{\beta}} + \lambda_{c,2} g^{\alpha\beta} \frac{\partial K}{\partial x^{\beta}} = 0. \tag{S19}$$

*Mass conservation equation.* – The mass conservation equation reads:

$$\frac{\partial \rho}{\partial t} + \nabla_{\alpha} \left(\rho v^{\alpha}\right) = 0. \tag{S20}$$

Using the Christoffel symbols, the above mass conservation equation further reads:

$$\frac{\partial \rho}{\partial t} = -\frac{\partial \left(\rho v^{\alpha}\right)}{\partial x^{\alpha}} - \rho v^{\omega} \Gamma^{\alpha}_{\alpha\omega}. \tag{S21}$$

To close the above dynamic equations (S16), (S19), and (S21), one needs the constitutive equation for the pressure. Following Ref. [1], we assume a dependence of the pressure $P$ on the density $\rho$:

$$P = P_0 \ln\left(\frac{\rho}{\rho_0}\right), \tag{S22}$$

where $P_0 > 0$ is a reference pressure and $\rho_0 > 0$ is a reference density.

*Discussion on the curvature-sensing free energy.* – Since $Q^{\alpha}_{\beta} = q(n^{\alpha} n_{\beta} - g^{\alpha}_{\beta}/2)$ with $q = \sqrt{2 Q^{\alpha}_{\beta} Q^{\beta}_{\alpha}}$ being the locally average nematic order parameter quantifying the locally average cell elongation magnitude, the curvature-sensing free energy Eq. (S12) can be re-expressed as,

$$F_C = \int \mathrm{d}S \left[\frac{1}{2} h_c q \left(C^{\alpha}_{\beta} n_{\alpha} n^{\beta} - \frac{1}{2} C^{\gamma}_{\gamma}\right)\right] \tag{S23}$$

Compared with Ref. [2], which describes a fully developed active nematic phase and thus assumes $q = 1$, here we allow the nematic order parameter $q$ to vary in space and time. This can reflect the different cell elongation magnitudes in space. Nevertheless, whether assuming $q = 1$ or allowing $q$ to vary, it will not affect the orientation feature of cells.

#### II. SOME TYPICAL CURVED SUBSTRATE SURFACES STUDIED

##### A. Hemicylindrical geometry

Here, we consider a hemicylindrical geometry, as shown in Supplementary Note Figure S1. We parameterize such a curved substrate surface by the coordinates $(s, y)$. With this parameterization, we have the surface metric tensor $g_{\alpha\beta} = \delta_{\alpha\beta}$ and the surface curvature tensor:

$$C_{ss} = C_{ss}(s) \quad , \quad C_{sy} = C_{ys} = C_{yy} = 0. \tag{S24}$$

In particular, we assume $C_{ss} = C_0 \cos(2\pi s/L_s)$ [2]. Depending on the curvature magnitude $C_0$, such a curvature profile can mimic different curved geometries, including the hemicylindrical surface.

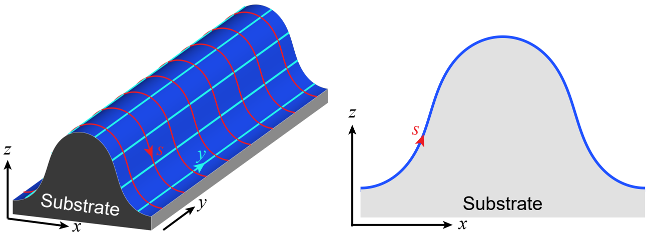


Supplementary Note Figure S1. Sketch of a hemicylindrical surface. (*Left*) 3D view, along with the $(s, y)$ coordinates. (*Right*) Cross-sectional view ($y =$ const).

###### 1. *Governing equations*

For the hemicylindrical geometry, we can simplify the governing equations as follows. First, the nematic tensor can be expressed as: $\boldsymbol{Q} = Q_{ss}\boldsymbol{e}_s \otimes \boldsymbol{e}_s + Q_{sy}\boldsymbol{e}_s \otimes \boldsymbol{e}_y + Q_{sy}\boldsymbol{e}_y \otimes \boldsymbol{e}_s - Q_{ss}\boldsymbol{e}_y \otimes \boldsymbol{e}_y$ with $\boldsymbol{e}_s$ and $\boldsymbol{e}_y$ being local, normalized base vector along the $s$ direction and the $y$ direction, respectively. $Q_{ss}$ and $Q_{sy}$ evolve according to:

$$\frac{\partial Q_{ss}}{\partial t} = -\left(v_s \frac{\partial Q_{ss}}{\partial s} + v_y \frac{\partial Q_{ss}}{\partial y}\right) + 2\Omega_{sy} Q_{sy} + \lambda E_{ss} + \Gamma H_{ss}, \tag{S25}$$

$$\frac{\partial Q_{sy}}{\partial t} = -\left(v_s \frac{\partial Q_{sy}}{\partial s} + v_y \frac{\partial Q_{sy}}{\partial y}\right) - 2\Omega_{sy} Q_{ss} + \lambda E_{sy} + \Gamma H_{sy}, \tag{S26}$$

where the strain rate tensor $E_{ss} = (\partial v_s/\partial s - \partial v_y/\partial y)/2$ and $E_{sy} = (\partial v_y/\partial s + \partial v_s/\partial y)/2$, the vorticity tensor $\Omega_{sy} = (\partial v_s/\partial y - \partial v_y/\partial s)/2$, and the molecular field tensor,

$$\begin{aligned} H_{ss} &= -A_1 Q_{ss} - \frac{1}{2} A_2 q^2 Q_{ss} + K_n \left(\frac{\partial^2 Q_{ss}}{\partial s^2} + \frac{\partial^2 Q_{ss}}{\partial y^2}\right) - \frac{1}{4}(h_{c,0} + h_{c,1} C_{ss}) C_{ss}, \\ H_{sy} &= -A_1 Q_{sy} - \frac{1}{2} A_2 q^2 Q_{sy} + K_n \left(\frac{\partial^2 Q_{sy}}{\partial s^2} + \frac{\partial^2 Q_{sy}}{\partial y^2}\right), \end{aligned} \tag{S27}$$

with $q = 2\sqrt{Q_{ss}^2 + Q_{sy}^2}$.

Second, the force balance equation reads:

$$\frac{\partial \sigma_{ss}}{\partial s} + \frac{\partial \sigma_{ys}}{\partial y} - \xi v_s + \lambda_{c,1} \frac{\partial C_{ss}}{\partial s} = 0, \tag{S28}$$

$$\frac{\partial \sigma_{sy}}{\partial s} + \frac{\partial \sigma_{yy}}{\partial y} - \xi v_y = 0, \tag{S29}$$

where the stress:

$$\begin{aligned}\sigma_{ss} &= -P + \eta\left(\frac{\partial v_s}{\partial s} - \frac{\partial v_y}{\partial y}\right) + \mathrm{X}\\ \sigma_{yy} &= -P + \eta\left(\frac{\partial v_y}{\partial y} - \frac{\partial v_s}{\partial s}\right) - \mathrm{X}\\ \sigma_{sy} &= \eta\left(\frac{\partial v_y}{\partial s} + \frac{\partial v_s}{\partial y}\right) + \mathrm{Y} + \mathrm{Z}\\ \sigma_{ys} &= \eta\left(\frac{\partial v_y}{\partial s} + \frac{\partial v_s}{\partial y}\right) + \mathrm{Y} - \mathrm{Z}\end{aligned} \tag{S30}$$

with

$$\mathrm{X} = -\lambda H_{ss} - \zeta Q_{ss} \quad , \quad \mathrm{Y} = -\lambda H_{sy} - \zeta Q_{sy} \quad , \quad \mathrm{Z} = 2H_{ss}Q_{sy} - 2Q_{ss}H_{sy}. \tag{S31}$$

Substituting Eq. (S30) into Eq. (S28) and (S29), we further obtain:

$$-\xi v_s + \eta\left(\frac{\partial^2 v_s}{\partial s^2} + \frac{\partial^2 v_s}{\partial y^2}\right) = \frac{\partial P}{\partial s} - \frac{\partial \mathrm{X}}{\partial s} - \frac{\partial \mathrm{Y}}{\partial y} + \frac{\partial \mathrm{Z}}{\partial y} - \lambda_{c,1}\frac{\partial C_{ss}}{\partial s}, \tag{S32}$$

$$-\xi v_y + \eta\left(\frac{\partial^2 v_y}{\partial s^2} + \frac{\partial^2 v_y}{\partial y^2}\right) = \frac{\partial P}{\partial y} + \frac{\partial \mathrm{X}}{\partial y} - \frac{\partial \mathrm{Y}}{\partial s} - \frac{\partial \mathrm{Z}}{\partial s}. \tag{S33}$$

Finally, the cell density $\rho$ evolves according to:

$$\frac{\partial \rho}{\partial t} = -\rho\left(\frac{\partial v_s}{\partial s} + \frac{\partial v_y}{\partial y}\right) - \left(v_s\frac{\partial \rho}{\partial s} + v_y\frac{\partial \rho}{\partial y}\right). \tag{S34}$$

##### 2. *Numerical simulation*

We perform numerical simulations of the full governing equations (S25), (S26), (S32), (S33), and (S34) in a rectangular domain of size $L_s \times L_y$ within the $(s, y)$ space. Considering the curved substrate used in our experiment where the hill-valley pattern appears alternately, we apply periodic boundary conditions in the $s$ direction. In addition, to mimic the hemicylindrical geometry, we also apply periodic boundary conditions along the $y$ direction and set $L_y$ much larger than the characteristic length scales of the active nematic system, including the hydrodynamic screening length $\ell_h = \sqrt{\eta/\xi}$, the nematic coherence length $\ell_n = \sqrt{K_n/A_2}$, and the nematic active length $\ell_a = \sqrt{K_n/|\zeta|}$.

We use the finite difference method to solve the nematic tensor field $Q_{ss}$ and $Q_{sy}$ and the cell density field $\rho$, where all the partial derivatives are calculated using the second-order central difference method. Specifically, we firstly update the nematic tensor field $Q_{ss}(\boldsymbol{r}, t) \to Q_{ss}(\boldsymbol{r}, t + \Delta t)$ and $Q_{sy}(\boldsymbol{r}, t) \to Q_{sy}(\boldsymbol{r}, t + \Delta t)$, and the cell density field $\rho(\boldsymbol{r}, t) \to \rho(\boldsymbol{r}, t + \Delta t)$, using a forward Euler scheme,

$$Q_{ss}(\boldsymbol{r}, t + \Delta t) = Q_{ss}(\boldsymbol{r}, t) + g_1(\boldsymbol{r}, t)\Delta t, \tag{S35}$$

$$Q_{sy}(\boldsymbol{r}, t + \Delta t) = Q_{sy}(\boldsymbol{r}, t) + g_2(\boldsymbol{r}, t)\Delta t, \tag{S36}$$

$$\rho(\boldsymbol{r}, t + \Delta t) = \rho(\boldsymbol{r}, t) + g_3(\boldsymbol{r}, t)\Delta t, \tag{S37}$$

where

$$g_1 = -\left(v_s\frac{\partial Q_{ss}}{\partial s} + v_y\frac{\partial Q_{ss}}{\partial y}\right) + 2\Omega_{sy}Q_{sy} + \lambda E_{ss} + \Gamma H_{ss}, \tag{S38}$$

$$g_2 = -v_s\frac{\partial Q_{sy}}{\partial s} - 2\Omega_{sy}Q_{ss} + \lambda E_{sy} + \Gamma H_{sy}, \tag{S39}$$

$$g_3 = -\left[\frac{\partial(\rho v_s)}{\partial s} + \frac{\partial(\rho v_y)}{\partial y}\right]. \tag{S40}$$

Next, based on the updated cell density field $\rho(\boldsymbol{r}, t+\Delta t)$, we update the pressure field as:

$$P(\boldsymbol{r}, t+\Delta t) = P_0 \ln\left[\frac{\rho(\boldsymbol{r}, t+\Delta t)}{\rho_0}\right]. \tag{S41}$$

Finally, based on the updated nematic tensor field $Q_{ss}(\boldsymbol{r}, t+\Delta t)$ and $Q_{sy}(\boldsymbol{r}, t+\Delta t)$, and the updated pressure field $P(\boldsymbol{r}, t+\Delta t)$, we employ the finite difference method to solve the force balance equations (S32) and (S33):

$$-\xi v_s(\boldsymbol{r}, t+\Delta t) + \eta\left(\frac{\partial^2 v_s(\boldsymbol{r}, t+\Delta t)}{\partial s^2} + \frac{\partial^2 v_s(\boldsymbol{r}, t+\Delta t)}{\partial y^2}\right) = g_4(\boldsymbol{r}, t+\Delta t), \tag{S42}$$

$$-\xi v_y(\boldsymbol{r}, t+\Delta t) + \eta\left(\frac{\partial^2 v_y(\boldsymbol{r}, t+\Delta t)}{\partial s^2} + \frac{\partial^2 v_y(\boldsymbol{r}, t+\Delta t)}{\partial y^2}\right) = g_5(\boldsymbol{r}, t+\Delta t), \tag{S43}$$

where

$$g_4 = \frac{\partial P}{\partial s} - \frac{\partial \mathrm{X}}{\partial s} - \frac{\partial \mathrm{Y}}{\partial y} + \frac{\partial \mathrm{Z}}{\partial y} - \lambda_{c,1}\frac{\partial C_{ss}}{\partial s}, \tag{S44}$$

$$g_5 = \frac{\partial P}{\partial y} + \frac{\partial \mathrm{X}}{\partial y} - \frac{\partial \mathrm{Y}}{\partial s} - \frac{\partial \mathrm{Z}}{\partial s}. \tag{S45}$$

Solving Eqs. (S42) and (S43) gives the updated flow field $v_s(\boldsymbol{r}, t+\Delta t)$ and $v_y(\boldsymbol{r}, t+\Delta t)$.

In our simulations, space and time steps are typically set as $\Delta s = \Delta y = 1/8$ and $\Delta t = 10^{-4}$, respectively. We set the initial nematic tensor field and the cell density field as small perturbations around the uniform equilibrium state. The initial velocity field $v_s(\boldsymbol{r}, t=0)$ and $v_y(\boldsymbol{r}, t=0)$ are calculated from the initial nematic tensor field $Q_{ss}(\boldsymbol{r}, t=0)$ and $Q_{sy}(\boldsymbol{r}, t=0)$, and the initial cell density field $\rho(\boldsymbol{r}, t=0)$ by solving the force balance equations.

We list the default parameters used in simulations in Table S1.

TABLE S1. List of other default parameter values used in simulations.

| Parameter | Description | Value |
|---|---|---|
| $\xi$ | Substrate friction | 0.2 |
| $\eta$ | Tissue shear viscosity | 1 |
| $P_0$ | Reference pressure | 10 |
| $\rho_0$ | Reference cell density | 1 |
| $\Gamma$ | Rotational viscosity | 1 |
| $\lambda$ | Flow alignment parameter | 0 |
| $A_1$ | Nematic elastic constant | $-0.1$ |
| $A_2$ | Nematic elastic constant | 0.2 |
| $K_n$ | Nematic elastic constant | 0.1 |
| $h_{c,0}$ | Constant curvature sensing parameter | 1 |
| $h_{c,1}$ | Mean curvature sensing parameter | 4 |
| $h_{c,2}$ | Gaussian curvature sensing parameter | 10 |
| $\zeta$ | Active stress | $-4$ |
| $\lambda_{c,1}$ | Mean curvotaxis traction | 10 |
| $\lambda_{c,2}$ | Gaussian curvotaxis traction | $-50$ |

##### 3. Theoretical analysis

*Simplified governing equations.* – Here, we consider a simple case where the system is uniform and invariant along the $y$ axis, i.e., $Q_{ss}=Q_{ss}(s)$, $Q_{sy}=Q_{sy}(s)$, $\rho=\rho(s)$, $v_s=v_s(s)$, and $v_y=v_y(s)$. Under such an assumption, the governing equations reduce to:

$$\frac{\partial Q_{ss}}{\partial t}=-v_s\frac{\partial Q_{ss}}{\partial s}+2\Omega_{sy}Q_{sy}+\lambda E_{ss}+\Gamma H_{ss}, \tag{S46}$$

$$\frac{\partial Q_{sy}}{\partial t}=-v_s\frac{\partial Q_{sy}}{\partial s}-2\Omega_{sy}Q_{ss}+\lambda E_{sy}+\Gamma H_{sy}, \tag{S47}$$

$$\frac{\partial \rho}{\partial t}=-\rho\frac{\partial v_s}{\partial s}-v_s\frac{\partial \rho}{\partial s}, \tag{S48}$$

$$-\xi v_s+\eta\frac{\partial^2 v_s}{\partial s^2}=\frac{\partial P}{\partial s}-\frac{\partial \mathrm{X}}{\partial s}-\lambda_{c,1}\frac{\partial C_{ss}}{\partial s}, \tag{S49}$$

$$-\xi v_y+\eta\frac{\partial^2 v_y}{\partial s^2}=-\frac{\partial \mathrm{Y}}{\partial s}-\frac{\partial \mathrm{Z}}{\partial s}, \tag{S50}$$

where the strain rate tensor, the vorticity tensor, and the molecular tensor now simplify to:

$$E_{ss}=\frac{1}{2}\frac{\partial v_s}{\partial s} \quad , \quad E_{sy}=\frac{1}{2}\frac{\partial v_y}{\partial s} \quad , \quad \Omega_{sy}=-\frac{1}{2}\frac{\partial v_y}{\partial s}, \tag{S51}$$

$$\begin{aligned} H_{ss}&=-A_1Q_{ss}-\frac{1}{2}A_2q^2Q_{ss}+K_n\frac{\partial^2 Q_{ss}}{\partial s^2}-\frac{1}{4}(h_{c,0}+h_{c,1}C_{ss})C_{ss},\\ H_{sy}&=-A_1Q_{sy}-\frac{1}{2}A_2q^2Q_{sy}+K_n\frac{\partial^2 Q_{sy}}{\partial s^2}. \end{aligned} \tag{S52}$$

*Non-flowing steady state.* – Now let us consider the non-flowing steady state solution, i.e., $v_s=v_y=0$, $\partial Q_{ss}/\partial t=0$, $\partial Q_{sy}/\partial t=0$, and $\partial\rho/\partial t=0$. Then, the steady state equations read:

$$H_{ss}=-A_1Q_{ss}-\frac{1}{2}A_2q^2Q_{ss}+K_n\frac{\partial^2 Q_{ss}}{\partial s^2}-\frac{1}{4}(h_{c,0}+h_{c,1}C_{ss})C_{ss}=0, \tag{S53}$$

$$H_{sy}=-A_1Q_{sy}-\frac{1}{2}A_2q^2Q_{sy}+K_n\frac{\partial^2 Q_{sy}}{\partial s^2}=0, \tag{S54}$$

$$\frac{\partial P}{\partial s}-\frac{\partial \mathrm{X}}{\partial s}-\lambda_{c,1}\frac{\partial C_{ss}}{\partial s}=0 \quad \Rightarrow \quad P=\mathrm{X}+\lambda_{c,1}C_{ss}+\text{const}, \tag{S55}$$

$$-\frac{\partial \mathrm{Y}}{\partial s}-\frac{\partial \mathrm{Z}}{\partial s}=0 \quad \Rightarrow \quad \mathrm{Y}+\mathrm{Z}=\text{const}, \tag{S56}$$

Note that at the steady-state, $\mathrm{X}=-\lambda H_{ss}-\zeta Q_{ss}=-\zeta Q_{ss}$, $\mathrm{Y}=-\lambda H_{sy}-\zeta Q_{sy}=-\zeta Q_{sy}$, and $\mathrm{Z}=2H_{ss}Q_{sy}-2H_{sy}Q_{ss}=0$. Thus, we further obtain:

$$P=-\zeta Q_{ss}+\lambda_{c,1}C_{ss}+\text{const}, \tag{S57}$$

$$Q_{sy}=0. \tag{S58}$$

Thus, $q=2|Q_{ss}|$. Substituting it into Eq. (S53) results in:

$$\frac{\partial^2 Q_{ss}}{\partial s^2}-\frac{A_1}{K_n}Q_{ss}-2\frac{A_2}{K_n}Q_{ss}^3-\frac{1}{4}\left(\frac{h_{c,0}}{K_n}+\frac{h_{c,1}}{K_n}C_{ss}\right)C_{ss}=0. \tag{S59}$$

We next try to get an analytical solution of Eq. (S59) in some limiting cases.

For a flat substrate surface, i.e., $C_{ss} = 0$, Eq. (S59) reduces to:

$$\frac{\partial^2 Q_{ss}}{\partial s^2} - \frac{A_1}{K_n} Q_{ss} - 2\frac{A_2}{K_n} Q_{ss}^3 = 0, \tag{S60}$$

which leads to two uniform solutions: $Q_{ss}^{(1)} = 0$ and $Q_{ss}^{(2)} = \pm Q_0^*$ with $Q_0^* = \sqrt{-A_1/(2A_2)} > 0$, corresponding to an isotropic phase and an nematic phase, respectively.

For a curved substrate surface, when the curvature is not too large, i.e., $|C_{ss}| \ll 1$, we can obtain an approximate solution for Eq. (S59). In such a limiting case, the solution $Q_{ss}(s)$ is close to the uniform solution for a flat substrate. We are interested in the nematic phase $Q_{ss} = -Q_0^* < 0$ and $Q_{sy} = 0$, which corresponds to a collective nematic orientation along the $y$ axis. Let us expand $Q_{ss}(s)$ to first-order terms in $C_{ss}$ around $Q_0 = -Q_0^*$:

$$Q_{ss}(s) = Q_0 + C_{ss}(s) Q_1(s) + O(C_{ss}^2). \tag{S61}$$

Substituting it into Eq. (S59) leads to differential equations for $Q_1(s)$:

$$\frac{\partial^2 Q_1}{\partial s^2} + 2\frac{A_1}{K_n} Q_1 - \frac{h_{c,0}}{4K_n} = 0, \tag{S62}$$

which results in:

$$Q_1(s) = D_1 \mathrm{e}^{\beta s} + D_2 \mathrm{e}^{-\beta s} + \frac{h_{c,0}}{8A_1}, \tag{S63}$$

with $\beta = \sqrt{-2A_1/K_n}$. Therefore, we obtain a first-order approximation of $Q_{ss}$ as:

$$Q_{ss} \approx -Q_0^* + C_{ss}\left(D_1 \mathrm{e}^{\beta s} + D_2 \mathrm{e}^{-\beta s} + \frac{h_{c,0}}{8A_1}\right). \tag{S64}$$

The coefficients $D_1$ and $D_2$ can be determined by boundary conditions. For example, if we assume a periodic boundary condition for $Q_{ss}$, i.e., $Q_{ss}(s=0) = Q_{ss}(s=L_s)$, we get $D_1 = D_2 = 0$. In such a case, we obtain a simple approximation for $Q_{ss}(s)$ as:

$$Q_{ss} \approx -Q_0^* + \frac{h_{c,0}}{8A_1} C_{ss}. \tag{S65}$$

Further, applying the pressure-density relation Eq. (S22) to Eq. (S57), we obtain an analytical expression for the profile of cell density at the steady-state:

$$\rho(s) = \rho_0 \exp\left[\frac{-\zeta Q_{ss}(s) + \lambda_{c,1} C_{ss}(s) + \mu_P}{P_0}\right] = \bar{\rho}_c \exp\left[\frac{-\zeta Q_{ss}(s) + \lambda_{c,1} C_{ss}(s)}{P_0}\right], \tag{S66}$$

where the unknown constants $\mu_P$ and $\bar{\rho}_c$ are to be determined by the mass conservation constraint $\int \rho \mathrm{d}s\mathrm{d}y = \rho_0 L_s L_y$. Substituting the approximation of $Q_{ss}$ (Eq. (S65)) into Eq. (S66), we further obtain:

$$\rho(s) \approx \rho_c \exp\left[\frac{\left(\lambda_{c,1} - \frac{h_{c,0}\zeta}{8A_1}\right) C_{ss}(s)}{P_0}\right], \tag{S67}$$

with $\rho_c = \bar{\rho}_c \exp(\zeta Q_0^*/P_0)$ a constant.

##### B. Hourglass geometry

Here, we consider an hourglass geometry, as shown in Supplementary Note Figure S2. We parameterize such a curved substrate surface by the coordinates $(s, \varphi)$.

Assume the hourglass is along the $x$-axis; thus, the curved surface is rotationally symmetric about the $x$-axis. Letting $C_{ss}(s)$ being the curvature along the $s$ direction, then

$$\frac{\mathrm{d}\vartheta}{\mathrm{d}s} = -C_{ss}(s), \tag{S68}$$

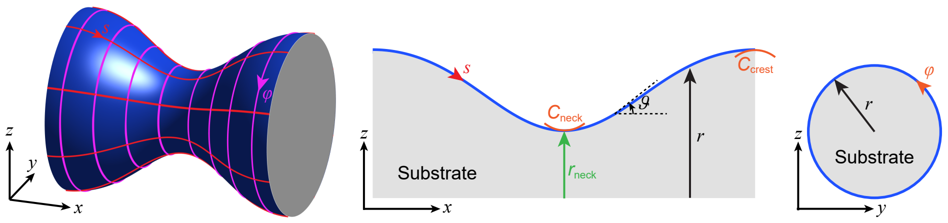


Supplementary Note Figure S2. Sketch of an hourglass substrate. (*Left*) 3D view, along with the $(s,\,\varphi)$ coordinates. (*Middle*) Cross-sectional view ($\varphi = \text{const}$), along with the definitions of $r_{\text{neck}}$, $C_{\text{neck}}$, and $C_{\text{crest}}$. (*Right*) Cross-sectional view ($s = \text{const}$).

where $\vartheta$ is the angle between the tangent of the $s$ arc and the $x$-axis (see Supplementary Note Figure S2). Thus,

$$\vartheta\left(s\right) = \vartheta_0 - \int_0^s C_{ss}\left(s\right)\mathrm{d}s, \tag{S69}$$

where $\vartheta_0 = \vartheta(s=0)$. The radius of the hourglass, i.e., the distance of the curved surface to the $x$-axis (Supplementary Note Figure S2), can be calculated by:

$$r\left(s\right) = r_0 + \int_0^s \sin\vartheta \mathrm{d}s, \tag{S70}$$

where $r_0 = r(s=0)$.

For such a surface, we can compute the basis vectors for the curvilinear coordinate system $(s,\varphi)$ as, $\boldsymbol{g}_s = \boldsymbol{e}_s$ and $\boldsymbol{g}_\varphi = r\boldsymbol{e}_\varphi$, with $\boldsymbol{e}_s$ and $\boldsymbol{e}_\varphi$ being the local, unit vectors along the $s$ direction and the $\varphi$ direction, respectively. Then, we can obtain the surface metric tensor,

$$(g_{\alpha\beta}) = (\boldsymbol{g}_\alpha \cdot \boldsymbol{g}_\beta) = \begin{pmatrix} g_{ss} & g_{s\varphi} \\ g_{\varphi s} & g_{\varphi\varphi} \end{pmatrix} = \begin{pmatrix} 1 & 0 \\ 0 & r^2 \end{pmatrix} \quad \Rightarrow \quad g = r^2 \tag{S71}$$

and the surface curvature tensor,

$$\left(C^\alpha_\beta\right) = \begin{pmatrix} C_{ss} & 0 \\ 0 & \dfrac{1}{r}\cos\vartheta \end{pmatrix} \tag{S72}$$

##### 1. *Governing equations*

Using the coordinates $(s,\,\varphi)$, the nematic tensor can be expressed as $\boldsymbol{Q} = Q^s_s \boldsymbol{g}_s \otimes \boldsymbol{g}^s + Q^s_\varphi \boldsymbol{g}_s \otimes \boldsymbol{g}^\varphi + Q^\varphi_s \boldsymbol{g}_\varphi \otimes \boldsymbol{g}^s + Q^\varphi_\varphi \boldsymbol{g}_\varphi \otimes \boldsymbol{g}^\varphi$ with $Q^\varphi_s = Q^s_\varphi / r^2$ and $Q^\varphi_\varphi = -Q^s_s$. Thus, the dynamic equation of the nematic tensor reads:

$$\frac{\partial Q^s_s}{\partial t} = -\left(v^s \frac{\partial Q^s_s}{\partial s} + v^\varphi \frac{\partial Q^s_s}{\partial \varphi}\right) + 2\frac{1}{r}\frac{\partial r}{\partial s} v^\varphi Q^s_\varphi + 2\Omega^\varphi_s Q^s_\varphi + \lambda E^s_s + \Gamma H^s_s, \tag{S73}$$

$$\frac{\partial Q^s_\varphi}{\partial t} = -\left(v^s \frac{\partial Q^s_\varphi}{\partial s} + v^\varphi \frac{\partial Q^s_\varphi}{\partial \varphi}\right) - 2r\frac{\partial r}{\partial s} v^\varphi Q^s_s + \frac{1}{r}\frac{\partial r}{\partial s} v^s Q^s_\varphi + 2\Omega^s_\varphi Q^s_s + \lambda E^s_\varphi + \Gamma H^s_\varphi, \tag{S74}$$

where

$$E^s_s = \frac{1}{2}\left(\frac{\partial v^s}{\partial s} - \frac{1}{r}\frac{\partial r}{\partial s} v^s - \frac{\partial v^\varphi}{\partial \varphi}\right) \quad , \quad E^s_\varphi = \frac{1}{2}\left(\frac{\partial v^s}{\partial \varphi} + r^2 \frac{\partial v^\varphi}{\partial s}\right), \tag{S75}$$

$$\Omega^s_\varphi = \frac{1}{2}\left(r^2 \frac{\partial v^\varphi}{\partial s} + 2r\frac{\partial r}{\partial s} v^\varphi - \frac{\partial v^s}{\partial \varphi}\right) \quad , \quad \Omega^\varphi_s = \frac{1}{2}\left(\frac{1}{r^2}\frac{\partial v^s}{\partial \varphi} - 2\frac{1}{r}\frac{\partial r}{\partial s} v^\varphi - \frac{\partial v^\varphi}{\partial s}\right) = -\frac{1}{r^2}\Omega^s_\varphi, \tag{S76}$$

$$
\begin{aligned}
H_s^s = & - A_1 Q_s^s - \frac{1}{2} A_2 q^2 Q_s^s - \frac{1}{4}[h_{c,0} + h_{c,1}(C_s^s + C_\varphi^\varphi) + h_{c,2} C_s^s C_\varphi^\varphi] \left(C_s^s - C_\varphi^\varphi\right) \\
& + K_n \left[ -4 \frac{1}{r^2} \left(\frac{\partial r}{\partial s}\right)^2 Q_s^s + \frac{\partial^2 Q_s^s}{\partial s^2} + \frac{1}{r}\frac{\partial r}{\partial s}\frac{\partial Q_s^s}{\partial s} + \frac{1}{r^2}\frac{\partial^2 Q_s^s}{\partial \varphi^2} - 4\frac{1}{r^3}\frac{\partial r}{\partial s}\frac{\partial Q_\varphi^s}{\partial \varphi} \right],
\end{aligned} \tag{S77}
$$

$$
H_\varphi^s = -A_1 Q_\varphi^s - \frac{1}{2} A_2 q^2 Q_\varphi^s + K_n \left[ 4\frac{1}{r}\frac{\partial r}{\partial s}\frac{\partial Q_s^s}{\partial \varphi} + \frac{\partial^2 Q_\varphi^s}{\partial s^2} - \frac{1}{r}\frac{\partial r}{\partial s}\frac{\partial Q_\varphi^s}{\partial s} + \frac{1}{r^2}\frac{\partial^2 Q_\varphi^s}{\partial \varphi^2} - \frac{1}{r}\frac{\partial^2 r}{\partial s^2} Q_\varphi^s - 3\frac{1}{r^2}\left(\frac{\partial r}{\partial s}\right)^2 Q_\varphi^s \right]. \tag{S78}
$$

The force balance equation reads:

$$
\frac{\partial \sigma_s^s}{\partial s} + \frac{1}{r^2}\frac{\partial \sigma_\varphi^s}{\partial \varphi} + \frac{1}{r}\frac{\partial r}{\partial s}\left(\sigma_s^s - \sigma_\varphi^\varphi\right) - \xi v^s + \lambda_{c,1}\frac{\partial\left(C_s^s + C_\varphi^\varphi\right)}{\partial s} + \lambda_{c,2}\frac{\partial\left(C_s^s C_\varphi^\varphi\right)}{\partial s} = 0, \tag{S79}
$$

$$
\frac{\partial \sigma_s^\varphi}{\partial s} + \frac{1}{r^2}\frac{\partial \sigma_\varphi^\varphi}{\partial \varphi} + 2\frac{1}{r}\frac{\partial r}{\partial s}\sigma_s^\varphi + \frac{1}{r^3}\frac{\partial r}{\partial s}\sigma_\varphi^s - \xi v^\varphi = 0, \tag{S80}
$$

where the stress:

$$
\begin{aligned}
\sigma_s^s &= -P + 2\eta E_s^s - \lambda H_s^s - \zeta Q_s^s \\
\sigma_\varphi^s &= 2\eta E_\varphi^s - \lambda H_\varphi^s - \zeta Q_\varphi^s + 2H_\varphi^s Q_s^s - 2H_s^s Q_\varphi^s \\
\sigma_s^\varphi &= 2\eta E_s^\varphi - \lambda H_s^\varphi - \zeta Q_s^\varphi - 2H_s^\varphi Q_s^s + 2H_s^s Q_s^\varphi \\
\sigma_\varphi^\varphi &= -P - 2\eta E_s^s + \lambda H_s^s + \zeta Q_s^s
\end{aligned} \tag{S81}
$$

Further substituting Eq. (S81) into Eqs. (S79) and (S80), we obtain:

$$
\begin{aligned}
& - \left[ \xi + \eta \frac{1}{r}\frac{\partial^2 r}{\partial s^2} + \eta\frac{1}{r^2}\left(\frac{\partial r}{\partial s}\right)^2 \right] v^s + \eta\frac{1}{r}\frac{\partial r}{\partial s}\frac{\partial v^s}{\partial s} + \eta\frac{\partial^2 v^s}{\partial s^2} + \eta\frac{1}{r^2}\frac{\partial^2 v^s}{\partial \varphi^2} - 2\eta\frac{1}{r}\frac{\partial r}{\partial s}\frac{\partial v^\varphi}{\partial \varphi} \\
& = \frac{\partial P}{\partial s} - \frac{\partial \mathrm{X}}{\partial s} - 2\frac{1}{r}\frac{\partial r}{\partial s}\mathrm{X} - \frac{1}{r^2}\frac{\partial \mathrm{Y}}{\partial \varphi} - \frac{1}{r^2}\frac{\partial \mathrm{Z}}{\partial \varphi} - \lambda_{c,1}\left(\frac{\partial C_s^s}{\partial s} + \frac{\partial C_\varphi^\varphi}{\partial s}\right) - \lambda_{c,2}\left(C_s^s\frac{\partial C_\varphi^\varphi}{\partial s} + C_\varphi^\varphi\frac{\partial C_s^s}{\partial s}\right),
\end{aligned} \tag{S82}
$$

$$
\begin{aligned}
& 2\eta\frac{1}{r^3}\frac{\partial r}{\partial s}\frac{\partial v^s}{\partial \varphi} - \xi v^\varphi + 3\eta\frac{1}{r}\frac{\partial r}{\partial s}\frac{\partial v^\varphi}{\partial s} + \eta\frac{\partial^2 v^\varphi}{\partial s^2} + \eta\frac{1}{r^2}\frac{\partial^2 v^\varphi}{\partial \varphi^2} \\
& = \frac{1}{r^2}\frac{\partial P}{\partial \varphi} + \frac{1}{r^2}\frac{\partial \mathrm{X}}{\partial \varphi} - \frac{1}{r^3}\frac{\partial r}{\partial s}\mathrm{Y} - \frac{1}{r^2}\frac{\partial \mathrm{Y}}{\partial s} - \frac{1}{r^3}\frac{\partial r}{\partial s}\mathrm{Z} + \frac{1}{r^2}\frac{\partial \mathrm{Z}}{\partial s},
\end{aligned} \tag{S83}
$$

where

$$
\mathrm{X} = -\lambda H_s^s - \zeta Q_s^s \quad , \quad \mathrm{Y} = -\lambda H_\varphi^s - \zeta Q_\varphi^s \quad , \quad \mathrm{Z} = 2H_\varphi^s Q_s^s - 2H_s^s Q_\varphi^s. \tag{S84}
$$

The mass conservation equation reads:

$$
\frac{\partial \rho}{\partial t} = -\left[\frac{\partial\left(\rho v^s\right)}{\partial s} + \frac{\partial\left(\rho v^\varphi\right)}{\partial \varphi}\right] - \frac{1}{r}\frac{\partial r}{\partial s}\rho v^s. \tag{S85}
$$

##### 2. *Numerical simulation*

We perform numerical simulations of the full governing equations (S73), (S74), (S85), (S82), and (S83) in a rectangular domain $[0, L_s] \times [0, 2\pi]$ within the $(s, \varphi)$ space. We apply periodic boundary conditions in both the $s$ direction and the $\varphi$ direction.

We use the finite difference method to solve the governing equations (S73), (S74), (S85), (S82), and (S83), similarly as demonstrated in Sec. II A 2.

We first apply the hourglass geometry to mimic the bead doublet in our experiments (Fig. 1(d)), by designing the curvature profile $C_{ss}(s)$ (equivalently, the tangent angle profile $\vartheta(s)$). Specifically, for a symmetrical bead doublet shape, we assume $\vartheta(s) = a_0 + a_1 s + a_2 s^2 + a_3 s^3$, thus $C_{ss}(s) = -a_1 - 2a_2 s - 3a_3 s^2$. The tangent angle profile $\vartheta(s)$ satisfies the boundary condition $\vartheta(s=0) = \vartheta(s=L_s/2) = 0$, and the curvature profile $C_{ss}$ satisfies the boundary condition $C_{ss}(s=0) = C_{\mathrm{neck}}$ and $C_{ss}(s=L_s/2) = C_{\mathrm{crest}}$, where $C_{\mathrm{neck}}$ and $C_{\mathrm{crest}}$ are the curvature at the neck region and the curvature at the crest region, respectively (along the $s$ direction), see Supplementary Note Figure S2. These boundary conditions result in the coefficients $a_0 = 0$, $a_1 = -C_{\mathrm{neck}}$, $a_2 = 2(2C_{\mathrm{neck}} + C_{\mathrm{crest}})/L_s$, and $a_3 = -4(C_{\mathrm{neck}} + C_{\mathrm{crest}})/L_s^2$. By varying the curvature at the neck region $C_{\mathrm{neck}}$ and the radius at the neck $r_{\mathrm{neck}}$, we can mimic the neck regions of different shapes and curvatures.

The default parameters used in simulations are listed in Table S1.

#### C. Dome-like geometry

Here, we consider a dome-like geometry, as shown in Supplementary Note Figure S3. We parameterize such a curved substrate surface by the coordinates $(s, \varphi)$; see Supplementary Note Figure S3(a,b).

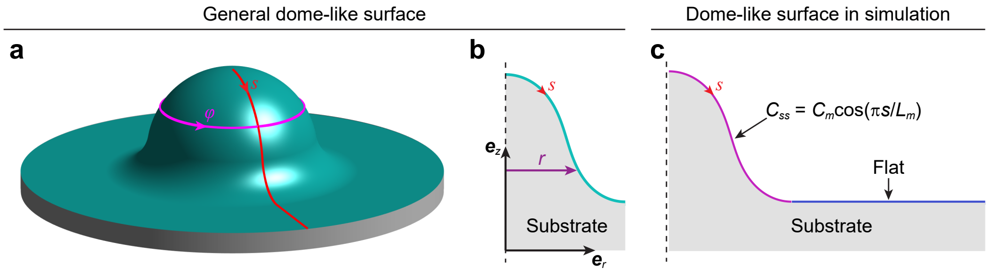


Supplementary Note Figure S3. Dome-like surface. (a, b) General case. (a) 3D view of a dome-like surface. Here, we also show the surface coordinates $(s, \varphi)$. (b) Cross-sectional view. (c) Dome-like surface geometry used in our simulation. It is composed of two regions, including a curved region and a flat plane of length $L_0$. The former, curved region is characterized by its curvature profile $C_{ss} = C_m \cos(\pi s/L_m)$ with $C_m$ quantifying the curvature magnitude and $L_m$ being the arc length.

Assume the dome-like surface is rotationally symmetric about the $z$-axis. Letting $C_{ss}(s)$ being the curvature along the $s$ direction, then

$$\frac{\mathrm{d}\vartheta}{\mathrm{d}s} = -C_{ss}(s), \tag{S86}$$

where $\vartheta$ is the angle between the tangent of the $s$ arc and the $r$ axis (Supplementary Note Figure S3(b)). Thus,

$$\vartheta(s) = \vartheta_0 - \int_0^s C_{ss}(s)\,\mathrm{d}s, \tag{S87}$$

where $\vartheta_0 = \vartheta(s=0)$. The radius, i.e., the distance of the curved surface to the $z$-axis (Supplementary Note Figure S3(b)), can be calculated by:

$$r(s) = r_0 + \int_0^s \cos\vartheta \mathrm{d}s, \tag{S88}$$

where $r_0 = r(s=0)$.

For such a surface, we can compute the basis vectors for the curvilinear coordinate system $(s, \varphi)$ as, $\boldsymbol{g}_s = \boldsymbol{e}_s$ and $\boldsymbol{g}_\varphi = r\boldsymbol{e}_\varphi$, with $\boldsymbol{e}_s$ and $\boldsymbol{e}_\varphi$ being the local, unit vectors along the $s$ direction and the $\varphi$ direction, respectively. Then, we can obtain the surface metric tensor,

$$(g_{\alpha\beta}) = (\boldsymbol{g}_\alpha \cdot \boldsymbol{g}_\beta) = \begin{pmatrix} g_{ss} & g_{s\varphi} \\ g_{\varphi s} & g_{\varphi\varphi} \end{pmatrix} = \begin{pmatrix} 1 & 0 \\ 0 & r^2 \end{pmatrix} \tag{S89}$$

and the surface curvature tensor,

$$(C^\alpha_\beta) = \begin{pmatrix} C_{ss} & 0 \\ 0 & -\dfrac{1}{r}\sin\vartheta \end{pmatrix} \tag{S90}$$

*1. Governing equations*

Using the coordinates $(s,\,\varphi)$, the nematic tensor can be expressed as $\boldsymbol{Q}=Q^s_s\boldsymbol{g}_s\otimes\boldsymbol{g}^s+Q^s_\varphi\boldsymbol{g}_s\otimes\boldsymbol{g}^\varphi+Q^\varphi_s\boldsymbol{g}_\varphi\otimes\boldsymbol{g}^s+Q^\varphi_\varphi\boldsymbol{g}_\varphi\otimes\boldsymbol{g}^\varphi$ with $Q^\varphi_s=Q^s_\varphi/r^2$ and $Q^\varphi_\varphi=-Q^s_s$. Thus, the dynamic equation of the nematic tensor reads:

$$\frac{\partial Q^s_s}{\partial t}=-\left(v^s\frac{\partial Q^s_s}{\partial s}+v^\varphi\frac{\partial Q^s_s}{\partial\varphi}\right)+2\frac{1}{r}\frac{\partial r}{\partial s}v^\varphi Q^s_\varphi+2\Omega^\varphi_s Q^s_\varphi+\lambda E^s_s+\Gamma H^s_s, \tag{S91}$$

$$\frac{\partial Q^s_\varphi}{\partial t}=-\left(v^s\frac{\partial Q^s_\varphi}{\partial s}+v^\varphi\frac{\partial Q^s_\varphi}{\partial\varphi}\right)-2r\frac{\partial r}{\partial s}v^\varphi Q^s_s+\frac{1}{r}\frac{\partial r}{\partial s}v^s Q^s_\varphi+2\Omega^s_\varphi Q^s_s+\lambda E^s_\varphi+\Gamma H^s_\varphi, \tag{S92}$$

where

$$E^s_s=\frac{1}{2}\left(\frac{\partial v^s}{\partial s}-\frac{1}{r}\frac{\partial r}{\partial s}v^s-\frac{\partial v^\varphi}{\partial\varphi}\right)\quad,\quad E^s_\varphi=\frac{1}{2}\left(\frac{\partial v^s}{\partial\varphi}+r^2\frac{\partial v^\varphi}{\partial s}\right), \tag{S93}$$

$$\Omega^s_\varphi=\frac{1}{2}\left(r^2\frac{\partial v^\varphi}{\partial s}+2r\frac{\partial r}{\partial s}v^\varphi-\frac{\partial v^s}{\partial\varphi}\right)\quad,\quad \Omega^\varphi_s=\frac{1}{2}\left(\frac{1}{r^2}\frac{\partial v^s}{\partial\varphi}-2\frac{1}{r}\frac{\partial r}{\partial s}v^\varphi-\frac{\partial v^\varphi}{\partial s}\right)=-\frac{1}{r^2}\Omega^s_\varphi, \tag{S94}$$

$$\begin{aligned}H^s_s=&-A_1Q^s_s-\frac{1}{2}A_2q^2Q^s_s-\frac{1}{4}[h_{c,0}+h_{c,1}(C^s_s+C^\varphi_\varphi)+h_{c,2}C^s_sC^\varphi_\varphi]\left(C^s_s-C^\varphi_\varphi\right)\\&+K_n\left[-4\frac{1}{r^2}\left(\frac{\partial r}{\partial s}\right)^2Q^s_s+\frac{\partial^2Q^s_s}{\partial s^2}+\frac{1}{r}\frac{\partial r}{\partial s}\frac{\partial Q^s_s}{\partial s}+\frac{1}{r^2}\frac{\partial^2Q^s_s}{\partial\varphi^2}-4\frac{1}{r^3}\frac{\partial r}{\partial s}\frac{\partial Q^s_\varphi}{\partial\varphi}\right],\end{aligned} \tag{S95}$$

$$H^s_\varphi=-A_1Q^s_\varphi-\frac{1}{2}A_2q^2Q^s_\varphi+K_n\left[4\frac{1}{r}\frac{\partial r}{\partial s}\frac{\partial Q^s_s}{\partial\varphi}+\frac{\partial^2Q^s_\varphi}{\partial s^2}-\frac{1}{r}\frac{\partial r}{\partial s}\frac{\partial Q^s_\varphi}{\partial s}+\frac{1}{r^2}\frac{\partial^2Q^s_\varphi}{\partial\varphi^2}-\frac{1}{r}\frac{\partial^2 r}{\partial s^2}Q^s_\varphi-3\frac{1}{r^2}\left(\frac{\partial r}{\partial s}\right)^2Q^s_\varphi\right]. \tag{S96}$$

The force balance equation reads:

$$\frac{\partial\sigma^s_s}{\partial s}+\frac{1}{r^2}\frac{\partial\sigma^s_\varphi}{\partial\varphi}+\frac{1}{r}\frac{\partial r}{\partial s}\left(\sigma^s_s-\sigma^\varphi_\varphi\right)-\xi v^s+\lambda_{c,1}\left(\frac{\partial C^s_s}{\partial s}+\frac{\partial C^\varphi_\varphi}{\partial s}\right)+\lambda_{c,2}\left(C^s_s\frac{\partial C^\varphi_\varphi}{\partial s}+C^\varphi_\varphi\frac{\partial C^s_s}{\partial s}\right)=0, \tag{S97}$$

$$\frac{\partial\sigma^\varphi_s}{\partial s}+\frac{1}{r^2}\frac{\partial\sigma^\varphi_\varphi}{\partial\varphi}+2\frac{1}{r}\frac{\partial r}{\partial s}\sigma^\varphi_s+\frac{1}{r^3}\frac{\partial r}{\partial s}\sigma^s_\varphi-\xi v^\varphi=0, \tag{S98}$$

where the stress:

$$\begin{aligned}\sigma^s_s&=-P+2\eta E^s_s-\lambda H^s_s-\zeta Q^s_s\\\sigma^s_\varphi&=2\eta E^s_\varphi-\lambda H^s_\varphi-\zeta Q^s_\varphi+2H^s_\varphi Q^s_s-2H^s_sQ^s_\varphi\\\sigma^\varphi_s&=2\eta E^\varphi_s-\lambda H^\varphi_s-\zeta Q^\varphi_s-2H^\varphi_sQ^s_s+2H^s_sQ^\varphi_s\\\sigma^\varphi_\varphi&=-P-2\eta E^s_s+\lambda H^s_s+\zeta Q^s_s\end{aligned} \tag{S99}$$

Further substituting Eq. (S99) into Eqs. (S97) and (S98), we obtain:

$$\begin{aligned}&-\left[\xi+\eta\frac{1}{r}\frac{\partial^2 r}{\partial s^2}+\eta\frac{1}{r^2}\left(\frac{\partial r}{\partial s}\right)^2\right]v^s+\eta\frac{1}{r}\frac{\partial r}{\partial s}\frac{\partial v^s}{\partial s}+\eta\frac{\partial^2v^s}{\partial s^2}+\eta\frac{1}{r^2}\frac{\partial^2v^s}{\partial\varphi^2}-2\eta\frac{1}{r}\frac{\partial r}{\partial s}\frac{\partial v^\varphi}{\partial\varphi}\\&=\frac{\partial P}{\partial s}-\frac{\partial \mathrm{X}}{\partial s}-2\frac{1}{r}\frac{\partial r}{\partial s}\mathrm{X}-\frac{1}{r^2}\frac{\partial \mathrm{Y}}{\partial\varphi}-\frac{1}{r^2}\frac{\partial \mathrm{Z}}{\partial\varphi}-\lambda_{c,1}\left(\frac{\partial C^s_s}{\partial s}+\frac{\partial C^\varphi_\varphi}{\partial s}\right)-\lambda_{c,2}\left(C^s_s\frac{\partial C^\varphi_\varphi}{\partial s}+C^\varphi_\varphi\frac{\partial C^s_s}{\partial s}\right),\end{aligned} \tag{S100}$$

$$\begin{aligned}&2\eta\frac{1}{r^3}\frac{\partial r}{\partial s}\frac{\partial v^s}{\partial\varphi}-\xi v^\varphi+3\eta\frac{1}{r}\frac{\partial r}{\partial s}\frac{\partial v^\varphi}{\partial s}+\eta\frac{\partial^2v^\varphi}{\partial s^2}+\eta\frac{1}{r^2}\frac{\partial^2v^\varphi}{\partial\varphi^2}\\&=\frac{1}{r^2}\frac{\partial P}{\partial\varphi}+\frac{1}{r^2}\frac{\partial \mathrm{X}}{\partial\varphi}-\frac{1}{r^3}\frac{\partial r}{\partial s}\mathrm{Y}-\frac{1}{r^2}\frac{\partial \mathrm{Y}}{\partial s}-\frac{1}{r^3}\frac{\partial r}{\partial s}\mathrm{Z}+\frac{1}{r^2}\frac{\partial \mathrm{Z}}{\partial s},\end{aligned} \tag{S101}$$

where

$$\mathrm{X} = -\lambda H_s^s - \zeta Q_s^s \quad , \quad \mathrm{Y} = -\lambda H_\varphi^s - \zeta Q_\varphi^s \quad , \quad \mathrm{Z} = 2H_\varphi^s Q_s^s - 2H_s^s Q_\varphi^s. \tag{S102}$$

The mass conservation equation reads:

$$\frac{\partial \rho}{\partial t} = -\left[\frac{\partial\left(\rho v^s\right)}{\partial s} + \frac{\partial\left(\rho v^\varphi\right)}{\partial \varphi}\right] - \frac{1}{r}\frac{\partial r}{\partial s}\rho v^s. \tag{S103}$$

##### 2. *Numerical simulation*

We perform numerical simulations of the full governing equations (S91), (S92), (S103), (S100), and (S101) in a rectangular domain $[0, L_s] \times [0, 2\pi]$ within the $(s,\ \varphi)$ space. The dome-like surface in our simulation is characterized by three geometric parameters $(C_m,\ L_m,\ L_0)$, see Supplementary Note Figure S3(c). By default, we set $C_m = 0.5$, $L_m = 8$, and $L_0 = 8$.

We apply periodic boundary conditions in the $\varphi$ direction. At the outer boundary $s = L_s$, we assume fixed boundary conditions, that is,

$$Q_s^s|_{s=L_s} = \frac{1}{2} q_B \cos 2\theta_B \quad , \quad Q_\varphi^s|_{s=L_s} = \frac{1}{2} r_B q_B \sin 2\theta_B \tag{S104}$$

$$v^s|_{s=L_s} = 0 \quad , \quad v^\varphi|_{s=L_s} = 0 \tag{S105}$$

where $r_B = r(s = L_s)$; $q_B$ is the cell elongation magnitude at the boundary $s = L_s$; $\theta_B$ is the orientation angle of cells with respect to the $s$-direction at the boundary $s = L_s$. In our simulations, we set $q_B = 0$. In addition, the flow field is smooth at the center $s = 0$, thus, $|v^s(0,\varphi)| < +\infty$ and $v^\varphi(0,\varphi) = 0$. Further, letting $s \to 0$, the force balance equations (S100) and (S101) lead to,

$$v^s(0,\varphi) = \frac{1}{2\eta} \frac{\left.\dfrac{\partial\left(\mathrm{Y}+\mathrm{Z}\right)}{\partial \varphi}\right|_{s=0}}{\left(\left.\dfrac{\partial r}{\partial s}\right|_{s=0}\right)^2}. \tag{S106}$$

We use the finite difference method to solve the governing equations (S91), (S92), (S103), (S100), and (S101), similarly as demonstrated in Sec. II A 2. The default parameters used in simulations are listed in Table S1.

---


[1] S. Yabunaka and P. Marcq, Physical Review E **96**, 022406 (2017).
[2] S. Bell, S.-Z. Lin, J.-F. Rupprecht, and J. Prost, Physical Review Letters **129**, 118001 (2022).

#### III. SUPPLEMENTARY FIGURES FOR THE MAIN TEXT

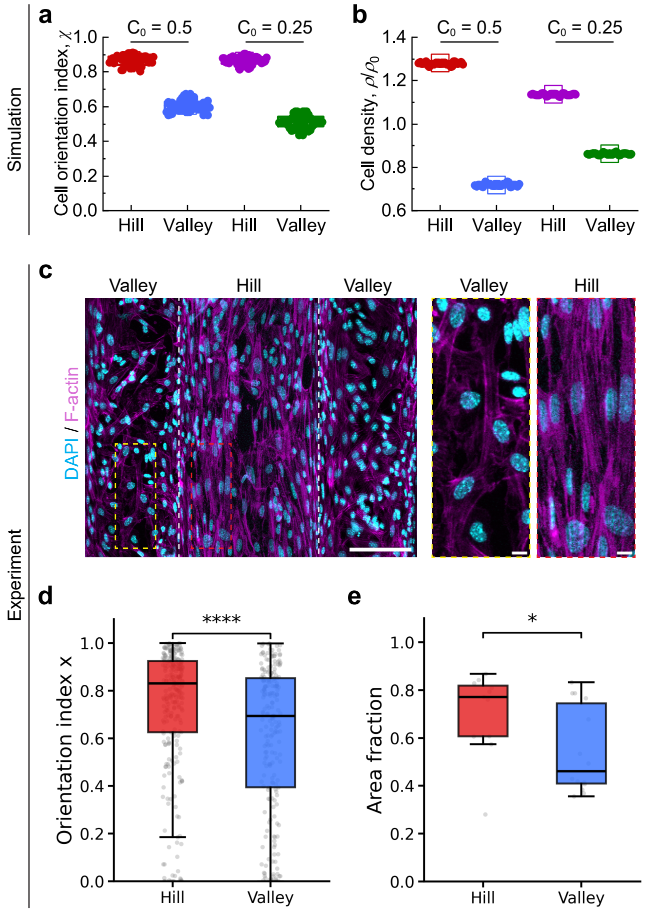


Supplementary Figure S1. Cell orientation and density profile on hemicylindrical geometries of different radius. (a, b) Numerical simulations. (a) Comparison of the cell orientation index $\chi$ for cells at the hill and valley regions, for two different curvature values: $C_0 = 0.5$, mimicking 100 $\mu$m hemicylinder, and $C_0 = 0.25$, mimicking 200 $\mu$m hemicylinder. (b) Comparison of the cell density $\rho$ for cells at the hill region and valley region, for two different curvature values: $C_0 = 0.5$, mimicking 100 $\mu$m hemicylinder and $C_0 = 0.25$, mimicking 200 $\mu$m hemicylinder. (c-e) Experiments. (c) Maximum projection of fluorescence images of TCs over a hemicylindrical substrate of 200 $\mu$m in radius. DAPI in cyan, F-actin in magenta. Scale bar, 100 $\mu$m. The dashed box represents the zoomed-in hill region (left) and valley region (right). Scale bar, 10 $\mu$m. (d) Quantification of TC orientations at hill and valley regions on 200 $\mu$m hemicylindrical substrates. Statistical significance was assessed using a two-sided Mann–Whitney U test. $^{*****}P < 0.0001$. $n = 273$ nuclei for hill, $n = 213$ nuclei for valley; 5 and 6 regions within the same substrates were analysed, respectively. $N = 3$ independent experiments. For comparison with 100 $\mu$m data, refer to Fig 1b, c. (e) Quantification of TC area fraction at hill and valley regions on 200 $\mu$m hemicylindrical substrates. Statistical significance was assessed using a two-sided Mann–Whitney U test. $^{*}P < 0.05$. $n = 11$ regions for hill and $n = 14$ regions for valley, from three independent experiments, were analysed respectively. For comparison with the 100 $\mu$m data, refer to Fig. 6(a, b) in the main text.

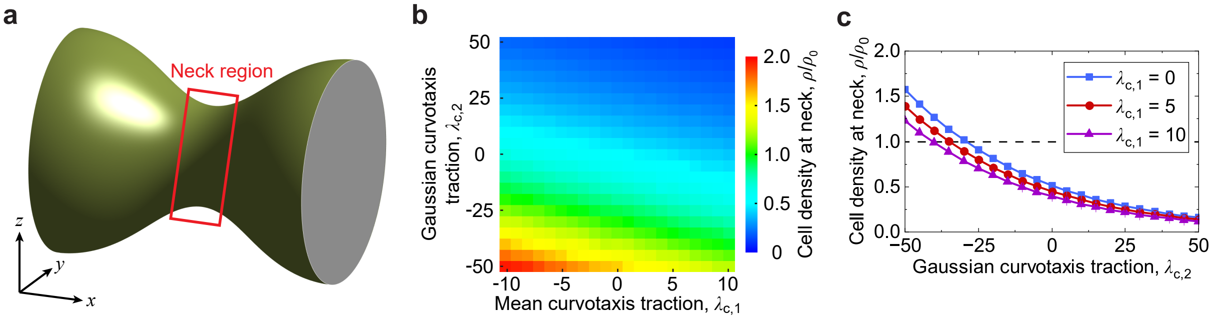


Supplementary Figure S2. Cell density on hourglass geometry. (a) Sketch of an hourglass substrate. (b) Phase diagram of the cell density at the neck region, regulated by the mean curvotaxis traction $\lambda_{c,1}$ and Gaussian curvotaxis traction $\lambda_{c,2}$. (c) The cell density at the neck region as a function of the Gaussian curvotaxis traction $\lambda_{c,2}$. See Table S1 for default parameter values used in simulations.

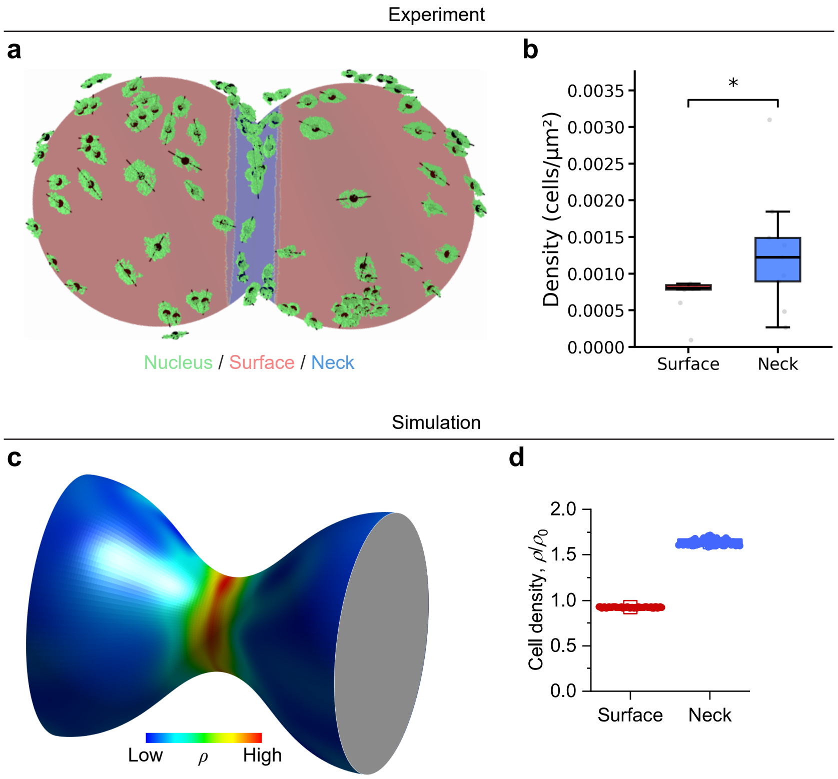


Supplementary Figure S3. Cell number density of theca cells on an hourglass substrate. (a, b) Experiment. (a) Segmentation of nuclei on the bead doublet is shown in green. Lines passing through the black dots, which indicate nuclear centroids, represent the nuclear long axis projected onto the local tangent plane of the spherical surface. The substrate is annotated to indicate the surface region (red) and the neck region (blue). (b) Comparison of TC density at the surface and neck regions on bead doublets. Statistical significance was assessed using a two-sided Mann–Whitney U test. $^*P < 0.05$; $n = 9$ bead doublets from 6 independent experiments were analyzed. (c, d) Simulation. (c) The cell density map obtained from simulations. (d) Simulation result of the cell density at the neck and surface regions. See Table S1 for default parameter values used in simulations.

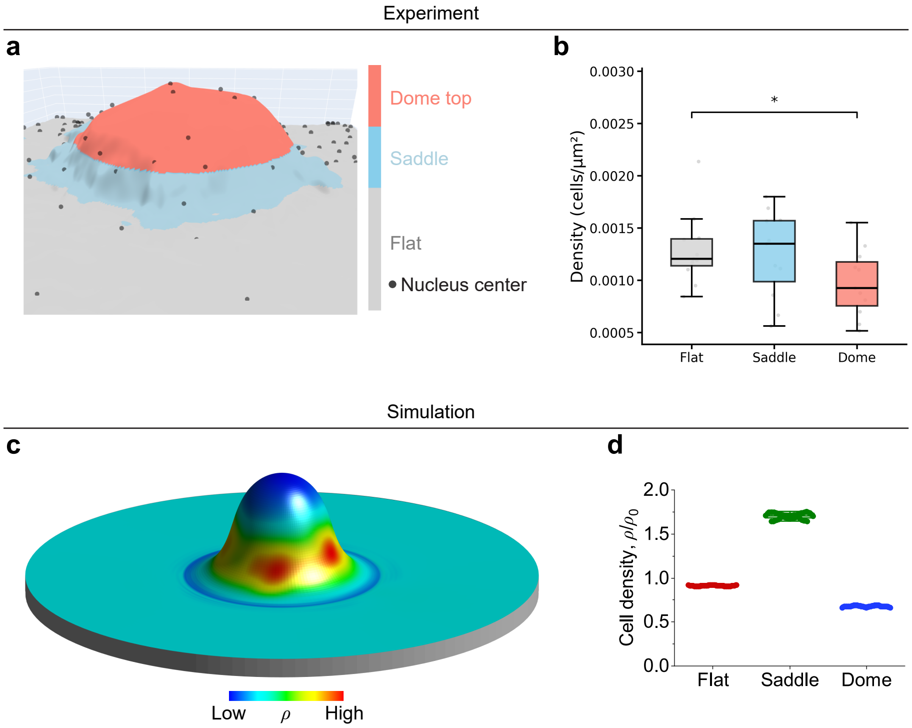


Supplementary Figure S4. Cell number density of theca cells on dome-like geometry: (a, b) experiment. (a) The substrate is annotated to indicate the dome surface region (red), the saddle region (blue), and the flat region (gray). The detected nuclei center is indicated by black dots. (b) Comparison of TC density at the flat region, saddle region, and dome surface on dome substrates. Statistical significance was assessed using a two-sided Mann–Whitney U test. $^{*}P < 0.05$ (Flat vs dome), n.s. (Saddle vs Dome), and n.s. (Flat vs Saddle); $n = 11$ dome substrates from 2 independent experiments were analyzed. (c, d) Simulation. (c) The cell density map obtained from simulations. (d) Simulation result of the cell density at the different regions. See Table S1 for default parameter values used in simulations.

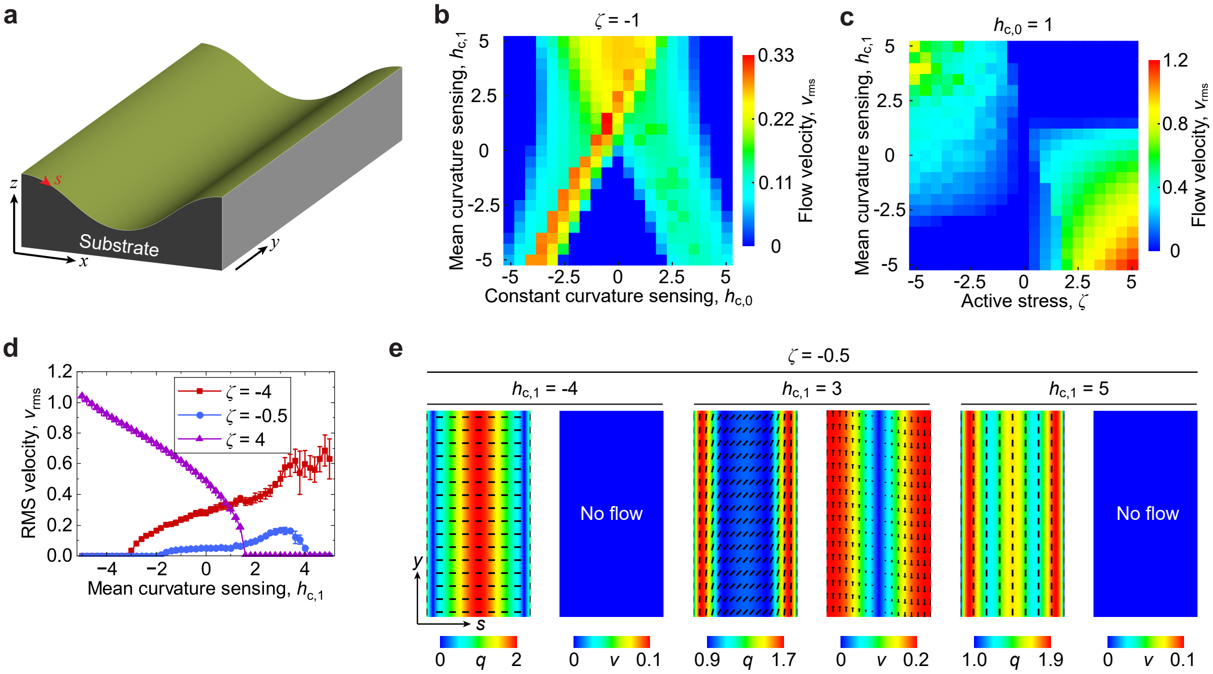


Supplementary Figure S5. Flows of a nematic cell layer over a hemicylindrical substrate. Here, we perform numerical simulations of Eqs. (S46) to (S50). (a) Schematic of a hemicylindrical substrate. The surface curvature is described by $C_{ss} = C_0 \cos(2\pi s/L_s)$. (b) The root-mean-square velocity $v_{\rm rms}$ regulated by the constant curvature sensing parameter $h_{c,0}$ and the mean curvature sensing parameter $h_{c,1}$, where $\zeta = -1$. (c) The root-mean-square velocity $v_{\rm rms}$ regulated by the active stress $\zeta$ and the mean curvature sensing parameter $h_{c,1}$, where $h_{c,0} = 1$. (d) The root-mean-square (RMS) velocity as a function of the mean curvature sensing parameter $h_{c,1}$ at different $\zeta$ values. (e) Typical cell orientation profiles and flow patterns at different $h_{c,1}$ values with $\zeta = -0.5$. Parameters: $r_{\rm neck} = 2$, $C_{\rm neck} = -0.5$, $C_{\rm crest} = 0.2$; see Table S1 for default parameter values used in simulations.

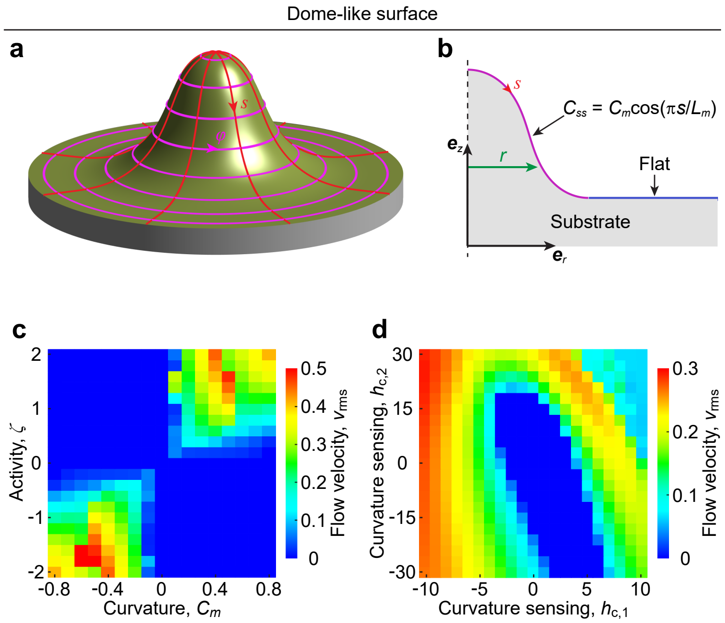


Supplementary Figure S6. Numerical simulation showing how substrate curvature, cell activity, and curvature-sensing parameters regulate the flow pattern of an active nematic layer on a dome-like surface. (a, b) Schematic of a dome-like curved substrate with a specific curvature profile. (a) 3D view of a dome-like surface with surface coordinates ($s$, $\varphi$). (b) Cross-sectional view of the dome-like surface geometry used in our simulation. It is composed of two regions, including a flat plane of length $L_0$ and a curved region with curvature profile $C_{ss} = C_m \cos(\pi s/L_m)$ where $C_m$ quantifies the curvature magnitude and $L_m$ is the total arc length. (c, d) Numerical simulation. (c) Phase diagram of the flow velocity $v_{\rm rms}$ regulated by the dome curvature $C_m$ and the activity $\zeta$. (d) Phase diagram of the flow velocity $v_{\rm rms}$ regulated by the curvature sensing parameters $h_{c,1}$ and $h_{c,2}$. Parameters: $L_0 = 8$, $L_m = 8$, $h_{c,0} = 1$, $\zeta = 1$, $\lambda_{c,1} = 0$, $\lambda_{c,2} = 0$; see Table S1 for other default parameter values.
